# Selection for seed size has indirectly shaped specialized metabolite abundance in oat (*Avena sativa* L.)

**DOI:** 10.1101/2021.08.18.454785

**Authors:** Lauren J. Brzozowski, Haixiao Hu, Malachy T. Campbell, Corey D. Broeckling, Melanie Caffe-Treml, Lucía Gutiérrez, Kevin P. Smith, Mark E. Sorrells, Michael A. Gore, Jean-Luc Jannink

**Author notes:** Correspondence: Lauren J. Brzozowski.

## Abstract

Plant breeding strategies to optimize metabolite profiles are necessary to develop health-promoting food crops. In oats (*Avena sativa* L.), seed metabolites are of interest for their antioxidant properties and their agronomic role in mitigating disease severity, yet have not been a direct target of selection in breeding. In a diverse oat germplasm panel spanning a century of breeding, we investigated the degree of variation of these specialized metabolites and how it has been molded by selection for other traits, like yield components. We also ask if these patterns of variation persist in modern breeding pools. Integrating genomic, transcriptomic, metabolomic and phenotypic analyses for three types of seed specialized metabolites – avenanthramides, avenacins, and avenacosides – we found reduced genetic variation in modern germplasm compared to diverse germplasm, in part due to increased seed size associated with more intensive breeding. Specifically, we found that abundance of avenanthramides increases with seed size, but additional variation is attributable to expression of biosynthetic enzymes, but avenacoside abundance decreases with seed size and plant breeding intensity. Overall, we show that increased seed size associated with plant breeding has uneven effects on the seed metabolome, and broadly contributes to understanding how selection shapes plant specialized metabolism.

## Introduction

Plants produce diverse arrays of specialized metabolites, generating a classification of hundreds of thousands of metabolites (Sorokina *et al*., 2021), that are nonessential for plant survival and frequently only found in specific plant lineages (Mutwil, 2020). Plant specialized metabolites are of interest for their role in biotic and abiotic stress tolerance as well as their implications for human health as nutraceutical compounds (Afendi *et al*., 2012; Jacobowitz & Weng, 2020). Plant breeding efforts to enhance specialized metabolite abundance in crop plants, however, are constrained by resource-intensive metabolomic phenotyping, genotype by environment interactions, and limited understanding of the genetic drivers of phenotypic variation in cultivated germplasm (Soltis & Kliebenstein, 2015). While advances in the study of model organisms like *Arabidopsis* have contributed to our understanding of specialized metabolism (Wager & Li, 2018), large scale studies on metabolomic diversity in cultivated germplasm – like glycoalkaloids in tomato (*Solanum lycopersicum* L.) (Zhu *et al*., 2018) and benzoxazinoids in maize (*Zea mays* L.) (Zhou *et al*., 2019) – provide information about specialized metabolism limited to specific lineages and in contexts more directly applicable for plant breeding programs. Overall, characterization of genomic variation and strategies to translate this information into widely applicable plant breeding strategies are critical steps to making specialized metabolite composition an accessible goal for plant breeding.

Studying specialized metabolites in cultivated plants in addition to wild progenitors or model organisms is important as specialized metabolite profiles may have also shifted in response to direct selection or indirect selection for other traits, or through genetic drift. While there is a longstanding prediction that cultivated plants would have reduced specialized metabolite concentration as compared to wild plants (as cultivated plants are more susceptible to biotic stress), there is not a consistent relationship between cultivation status and specialized metabolites across multiple species (Whitehead *et al*., 2017). Instead, differences in specialized metabolite abundance are frequently observed in distinct breeding pools and pedigrees. For instance, divergence in volatiles of roots has been noted in maize (Rasmann *et al*., 2005), and leaves in cranberry (Rodriguez-Saona *et al*., 2011) and there is variation in leaf glucosinolates in cultivated Brassicas (Poelman *et al*., 2008). For plant breeders, insight into how selection processes affected specialized metabolites can provide a basis for ongoing work and germplasm selection for breeding efforts.

We explored existing variation of specialized metabolites in oats (*Avena sativa* L.) and how the metabolomic profile has been shaped by plant breeding. Oats were domesticated from weedy progenitors (Loskutov, 2008) and, like other cereal crops, domesticated oats have increased seed size compared to wild species (Preece *et al*., 2017). Oats are used as livestock feed and have been an important part of human diet in some parts of Europe since before the Renaissance (Murphy & Hoffman, 1992). The nutraceutical benefits of fiber, skin soothing and general health promotion of oats were also noted in the first century CE by Dioscorides (Murphy & Hoffman, 1992). Today, oats are still known as a healthy whole grain (Singh *et al*., 2013; Stewart & McDougall, 2014), with high concentrations of unsaturated fats (Carlson *et al*., 2019) and heart health-promoting ß-glucans (Newell *et al*., 2012). Both have been the subject of plant breeding efforts, but yield and disease resistance are still predominant traits of interest for plant breeding (Haikka *et al*., 2020; González-Barrios *et al*., 2021). In addition to these health-promoting compounds, oat seeds contain multiple specialized metabolites (Sang & Chu, 2017) but, to the best of our knowledge, these metabolites have not been a direct target of selection. With this history, we predict that oat specialized metabolites may have been subject to genetic drift or indirect selection processes (e.g., for seed traits or disease resistance) leading to changes in patterns of variation. Characterizing the genetic bases of variation will provide a starting point for plant breeding.

We focused on three types of specialized metabolites in oat seed: avenanthramides, and the saponins avenacins and avenacosides. Avenanthramides are in highest concentration in the outer layers of the seed, most notably the aleurone layer (Liu & Wise, 2021), while the saponin avenacosides are concentrated in the endosperm (Önning *et al*., 1993). Avenanthramides have antioxidant properties (Meydani, 2009; Sang & Chu, 2017) that are retained through processing of oats into many consumer products (Pridal *et al*., 2018). The committed enzymes of avenanthramide biosynthesis have been characterized, and it is well-established that avenanthramides are the result of condensation between phenolic acids and anthranilic acid, products of different branches of aromatic amino acid biosynthesis (Collins, 2011; Wise, 2014; Li *et al*., 2019). Avenanthramides are associated with resistance to crown rust (pathogen *Puccinia coronata* f. sp. *avenae*), (Wise *et al*., 2008; Wise, 2014), and demonstrate variation in response to the environment (Emmons & Peterson, 2001; Peterson *et al*., 2005; Michels *et al*., 2020). The avenacins and avenacosides are both saponins that have been implicated in reducing plant fungal infections and in lowering cholesterol when consumed, but have received less attention for research and breeding (Sang & Chu, 2017). Core biosynthetic genes for avenacin biosynthesis have been identified in roots of the non-cultivated species, *Avena strigosa* (Kemen *et al*., 2014; Leveau *et al*., 2019), but whether variation in expression of these genes affects abundance in seed tissues of cultivated oat remains unknown.

Knowledge of biochemical pathways is a crucial foundation but, for plant breeding, it is important to further investigate whether variants that affect enzyme activity, or regulation, or pathway flux, or metabolite transport contribute to the observed phenotypic variation (Soltis & Kliebenstein, 2015). While loss of function mutations in biosynthetic enzymes are observed and employed by breeders for specialized metabolites in some crops (e.g., *Pun1* mutation prevents capsaicin production in pepper (Stewart *et al*., 2005)), mutations in regulatory elements are critical in others (e.g., transcription factor *Bt* mediates cucurbitacin accumulation in cucumber (Shang *et al*., 2014)). For oats, there is experimental evidence that avenanthramides increase in response to activation of systemic acquired resistance (salicylic acid mediated defense) (Wise, 2011, 2017; Wise *et al*., 2016), and degree of induction varies between oat genotypes (Wise *et al*., 2016), suggesting that regulatory variants could be an important target for selection. While expression of key biosynthetic enzymes has been profiled (Dimberg & Peterson, 2009; Wise, 2017), there has not been a genome-wide or transcriptome-wide association study to identify novel genes. We are not aware of comparable studies of saponins. In other crops, integrated genomic, transcriptomic and metabolomic analyses have been critical in understanding metabolic profiles. For instance, concomitant changes in fruit metabolome and fruit size have been characterized in tomatoes (Zhu *et al*., 2018).

We sought to integrate oat seed metabolomic, transcriptomic and genomic data to characterize genetic variation contributing to specialized metabolite abundance in oat seed. We also measured oat seed size to evaluate if selection on that yield component has affected specialized metabolite profiles. Using a diverse germplasm panel that includes oat varieties developed beginning in 1920 and an elite germplasm panel, we measured whole seed metabolome phenotypes and seed size and weight traits. In the diverse germplasm panel, we also conducted transcriptome sequencing of developing seed. We hypothesized that variation is greater in the diversity than the elite panel, and examined the relationship between seed traits and specialized metabolites in both of these panels. We also investigated the relative roles of variation in regulation and known biosynthetic enzyme pathway genes in mediating metabolite variance. To test these predictions, we conducted a genome-wide and a transcriptome-wide association study (GWAS, TWAS, respectively) and eQTL analysis for metabolites and seed traits. Overall, this work provides insight into breeding for oat specialized metabolites and more broadly adds to our foundation of how the relative contributions of genetic variation in regulation versus direct biosynthesis shapes phenotypic variation of specialized metabolites in crop plants.

## Materials and methods

### Oat germplasm

We used two germplasm panels of inbred lines, a diversity panel intended to capture genetic diversity in cultivated oats and an elite panel consisting of lines selected from the North American uniform oat performance nursery. These germplasm panels have been previously described in Campbell *et al*. (2021) and Hu *et al*. (2021). In the diversity panel, there were 368 entry genotypes (inbred lines) and seven check genotypes planted in an augmented design at Ithaca, New York, US in 2018. Six genotypes that lacked both genotyping data and gene expression data were removed from our analysis. The elite panel consisted of inbred lines, and was evaluated in three northern US environments (Minnesota, “MN”; South Dakota, “SD”; Wisconsin, “WI”) in 2017 in an augmented design with 232 entries and three checks. Nineteen entries were included in both the diversity and elite panels, and were removed from the elite panel analyses to compare independent sets of germplasm.

### Oat seed secondary metabolite phenotypes

We profiled the seed metabolome in the oat diversity panel and elite panel. Detailed descriptions of extraction and processing of these samples has been previously (Campbell *et al*., 2021; Hu *et al*., 2021) and is provided here in **Method S1**. Briefly, extractions and measurements were conducted at the Bioanalysis and Omics Center of the Analytical Resources Core (“ARC-BIO”), at Colorado State University (Fort Collins, CO, USA). Briefly, 50 seeds were dehulled, homogenized and extracted using a biphasic extraction method to separate polar and non-polar compounds. Chromatography analysis of the polar compounds (aqueous layer) was was done using a Waters Acquity UPLC system with a Waters Acquity UPLC CSH Phenyl Hexyl column (1.7 μM, 1.0 x 100 mm) and a Waters Xevo G2 TOF-MS with an electrospray source in positive mode. Mass features were annotated by first searching against an in-house spectra and retention time database using RAMSearch (Broeckling *et al*., 2016) and then by using MSFinder (Tsugawa *et al*., 2016). Names and spectra of the specialized metabolites are given in **Table S1**. The mass spectra of the specialized metabolites were positively annotated by these methods in the diversity panel, which was analyzed in 2018. Many of the specialized metabolites were also annotated in the elite panel (measured in 2017), and missing annotations were completed by comparing spectra to the diversity panel and published mass spectra for avenanthramides (de Bruijn *et al*., 2019), avenacins (Leveau *et al*., 2019) and avenacosides (Bahraminejad *et al*., 2008). The final phenotype reported was the relative signal intensity (relative concentration) of each metabolite.

Best linear unbiased predictions (BLUPs) were calculated for each metabolite for the diversity panel, and separately for each environment of the elite panel. To account for skew, data were log2 transformed. Then, relative concentration of each metabolite was modeled with a linear mixed model in R (R Core Team, 2016) with lme4 (Bates *et al*., 2015). For each metabolite, there were fixed effects of whether the genotype was a replicated check and days to heading (“DTH”) as a numeric covariate, and random effects of experimental block, batch in which the sample was run on the LCMS, and genotype. Outliers were defined as having studentized residual >3, and were removed, and the model was recalculated. Effect significance of the DTH covariate is shown in (**Table S2**). The BLUPs were then deregressed (Garrick *et al*., 2009). The deregressed BLUPs (drBLUPs) were used in all following analyses. Pearson’s correlations were estimated between phenotypes using the ‘cor.test’ function in R.

### Oat seed size and mass phenotypes

After dehulling, 50 seeds were used for evaluating seed length, width and height. The seeds were scanned with a two-dimensional scanner, where seed length and width were extracted with the software WinSeedle (Regent Instrument Canada Inc., version 2017). Seed height was measured separately using an electronic caliper manually with accuracy of 0.01mm. Seed length and width measurements are not available from the elite panel that was evaluated in South Dakota. Seed volume was estimated as an ellipsoid (Clohessy *et al*., 2018), and surface area of an ellipsoid was estimated by S≈ 4π *((lw)1.6+(lh)1.6+(wh)1.6)) / 3)^ (1/1.6). Separately, 100 hand dehulled seeds (hundred kernel weight, “HKW”) and their respective hulls (“HHW”) were weighed and the percent groat (kernel) was calculated as the percent of total weight (kernel plus hull weight). Deregressed BLUPs were then calculated from untransformed values in the same manner as the metabolites (above), and used in all further analysis. The relationship between drBLUPs of seed traits and metabolites was modeled with a linear model and effect significance was tested by ANOVA.

### Oat variety release year

We conducted an extensive literature search to determine the year of variety release for as many varieties in the diversity panel as possible. Most varieties were identified from information on USDA GRIN (https://npgsweb.ars-grin.gov), some in Triticeae Toolbox (https://triticeaetoolbox.org/POOL), others in US (https://apps.ams.usda.gov/), Canada (https://www.inspection.gc.ca/english/plaveg/pbrpov/cropreport/oat) or Europe (https://ec.europa.eu/food/plant/plant_propagation_material/plant_variety_catalogues_databases/) plant registrations, and finally as published variety releases. In sum, we identified the year of variety release for 155 varieties (**Table S3**).

### Genotyping and genome-wide association study

Genotyping-by-sequencing data was retrieved from T3/Oat (https://oat.triticeaetoolbox.org/), filtered to remove markers with more than 60% missingness and markers with a minor allele frequency of less than 0.02, and then imputed using the glmnet function (Friedman *et al*., 2010) in R. Overall, there were 73,527 markers, of which 54,284 could be anchored to the genome (PepsiCO OT3098v1; https://wheat.pw.usda.gov/GG3/graingenes_downloads/oat-ot3098-pepsico). All 54,284 SNPs were used for the diversity panel, and 54,219 SNPs were used for the elite panel after these imputed SNPs were again filtered by minor allele frequency. Kinship matrices were calculated for the diversity and elite panels with their SNPs using the A.mat function, and genomic heritability (de los Campos *et al*., 2015) was calculated using the kin.blup function in rrBLUP (Endelman, 2011). Genetic correlations were calculated in sommer using the mmer and cov2cor functions (Covarrubias-Pazaran, 2016). Principal components to use as covariates to account for population structure were calculated using the prcomp function in R. The first 25 PCs were calculated, and the scree plot was visually examined to determine the number of PCs to use in future analyses (**Figure S1**). Five PCs were chosen for the diversity panel and 4 PCs were chosen for the elite panel. Genome-wide association study (GWAS) was conducted for each phenotype (drBLUP) in statgenGWAS (Rossum & Kruijer, 2020) using the PCs as covariates and the kinship matrix. For GWAS results, *P*-values were adjusted with a bonferroni correction on a per-trait basis and SNPs with a *pBonf* < 0.05 were considered significant. To determine if any results colocalized with known QTL for crown rust, crown rust QTL were recorded from recent publications and mapped to the latest genome version (**Table S4**) (Lin *et al*., 2014; Babiker *et al*., 2015; McNish *et al*., 2020; Zhao *et al*., 2020).

### Transcriptome analyses of oat diversity panel

Developing oat seed tissue was dissected, and RNA was extracted using a hot borate protocol at 23 DAA as this time point showed slightly higher correlation between transcript and relative concentration of metabolites than other sampled developmental time points (Hu *et al*., 2020). RNAseq reads were aligned to the oat transcriptome using Salmon v0.12 (Patro *et al*., 2017) and transformed using variance stabilizing transformation in DESeq2 (Love *et al*., 2014) as described by Hu *et al*., (2020). For these analyses, we removed all transcripts expressed in fewer than 50% of samples as these are not useful for TWAS, leaving 54% of the original set (29,385). We examined the median absolute deviance of these transcripts to look for outliers and none exceeded a cutoff of MAD>10. Deregressed BLUPs were then calculated in sommer (Covarrubias-Pazaran, 2016) using the ‘mmer’ function. For each transcript, there were fixed effects of whether the genotype was a replicated check, the plate in which RNA was extracted from, and days to heading (“DTH”) as a numeric covariate, and random effects of experimental block and genotype. In all, 22,638 transcripts had converged drBLUPs (non-zero heritability). To remove any additional factors associated with experimental design, we ran probabilistic estimation of expression residuals (PEER) and found that k=5 factors (determined by visual examination of scree plot) was sufficient (**Figure S2**).

We then conducted a transcriptome wide association study (TWAS) and enrichment analyses. We used the transcript PEER residuals and a kinship matrix, as well as five genomic PCs as covariates for TWAS on the metabolite and seed trait drBLUPs. We implemented TWAS using the ‘createGData’ and ‘runSingleTraitGwas’ functions in the statgenGWAS package (Rossum & Kruijer, 2020). *P*-values were adjusted per trait using a false discovery rate adjustment, and transcripts with *p_FDR_* < 0.05 were considered significant. The adjusted *p*-values for all transcripts were used in gene ontology (GO) enrichment analysis for each of the phenotypes for biological processes GO terms. Enrichment analysis was implemented in the R package topGO, where significance was determined based on the default “weight01” algorithm followed by a Fisher test (Alexa & Rahnenfuhrer, 2016). Finally, transcripts had previously been assigned to temporally covarying groups (Hu *et al*., 2020) and these annotations were used to assign transcripts by date (8, 13, or 18 DAA) and direction (up or down) that expression pattern shifted. Those that changed on multiple dates were split into the two respective days. We tested for enrichment of any temporal and direction class using a hypergeometric test with the ‘phyper’ function in R.

We also identified transcripts associated with the avenanthramide biosynthetic pathway (beginning at PAL) and the preceding shikimate pathway using Ensemble Enzyme Prediction Pipeline (E2P2) annotations (Chae *et al*., 2014) of transcripts (**Table S5**).

### eQTL analysis

We implemented eQTL analysis in Matrix eQTL (Shabalin, 2012) in R using the PEER residuals for transcript counts and with five genomic PCs as covariates. SNPs were defined as significant eQTL at a threshold of *p_FDR_* < 0.05 per transcript. As only half of the transcripts are mapped, we did not differentiate between *cis* and *trans* eQTL, although future genome and transcriptome assemblies will facilitate this analysis.

## Results

### Heritability and correlations of specialized metabolites in oat seed

Specialized metabolites (avenanthramides, “AVNs”; avenacins, “AECs”; avenacosides, “AOSs”) were measured in seeds of a diverse germplasm panel evaluated in one environment and an elite set of oat germplasm evaluated in three environments. Genomic heritability was low to moderate for most metabolites, and some metabolites had heritability less than 0.05 (**Figure 1**). In general, there was a strong degree of phenotypic and genetic correlation within metabolite groups (e.g., within AVNs) across populations and environments, with the exception of avenacoside B (AOS_B) (**Figure 1**). In the diversity panel, the phenolic AVNs tended to have negative phenotypic and genetic correlations with both saponins (AEC and AOS), while AECs and AOSs were positively correlated (**Figure 1a**). This trend was less pronounced in the elite population phenotypes in most environments (**Figure 1b-d**). While there was still strong within-group correlation, there were no significant negative phenotypic correlations between phenolics and saponins.

**Figure 1.**
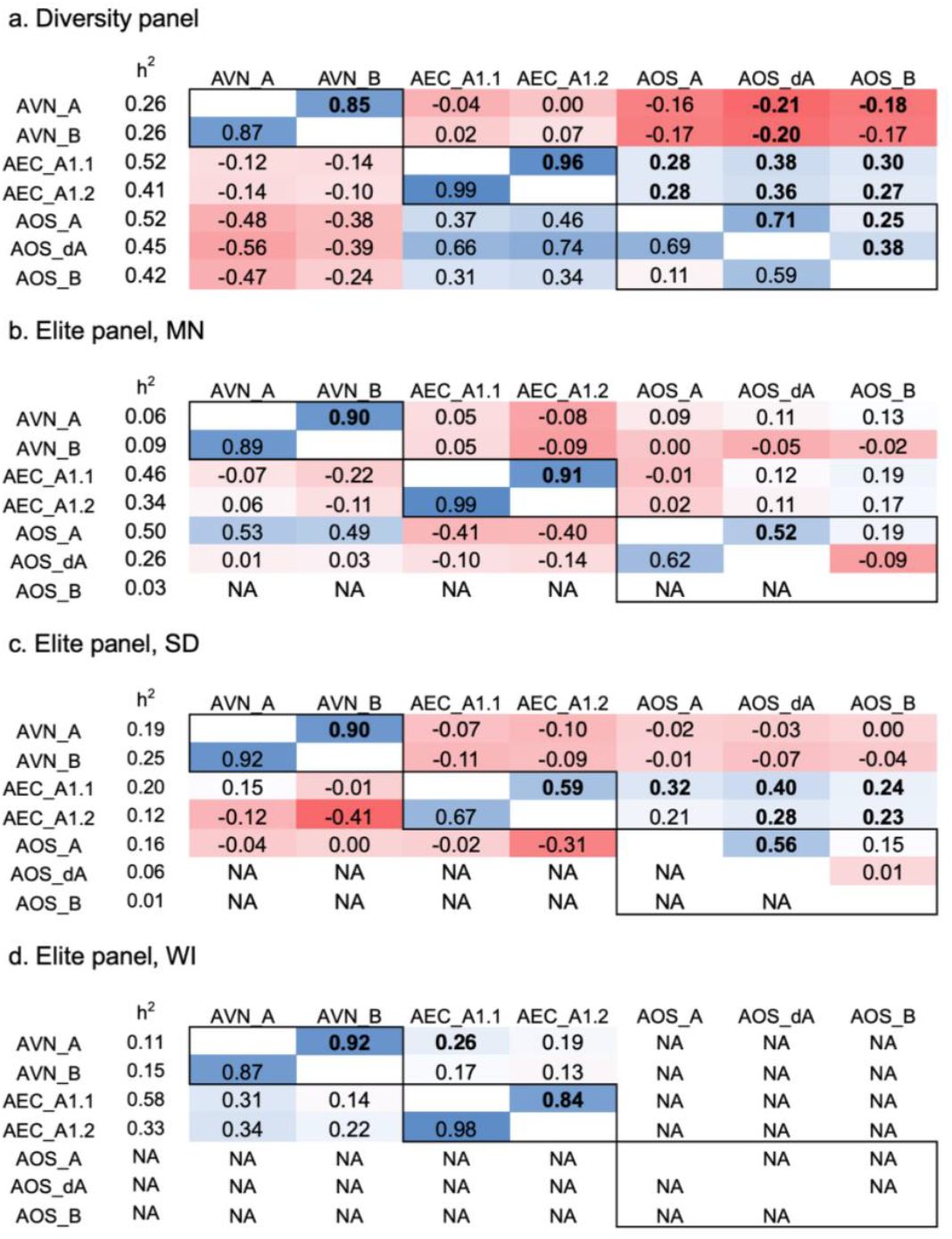
Phenotypic and genetic correlation of specialized metabolites in oat seed (avenanthramides, “AVN”; avenacins, “AEC”; avenacosides, “AOS”) in the (a) diverse panel evaluated only in New York, and elite panel evaluated in (b) Minnesota (“MN”), (c) South Dakota (“SD”), (d) Wisconsin (“WI”), USA. The specific type of metabolite is described in **Table S1**. The values in the top diagonal are Pearson’s phenotypic correlations, where bold indicates significance at the Bonferroni cutoff (*p*<0.001), the values in the bottom diagonal are genetic correlations with no associated statistical values, and h^2^ is the genomic heritability.

### Relationship between seed traits and specialized metabolites

We examined seed size traits in dehulled seeds (volume, surface area and surface area to volume ratio), as well as kernel and hull weight and percent groat (kernel). In general, heritability of the seed traits was greater than those of the specialized metabolites (**Table S6**) and seed volume was used for further analyses (diversity panel h^2^=0.72; elite panel Minnesota h^2^=0.50; elite panel Wisconsin h^2^=0.33). There were significant relationships between some metabolites and seed size (**Figure 2**; **Table S7**) and seed weight and composition (**Table S8**). In both the diversity and elite panel, relative concentration of AVNs (present in outer seed layers) increased with seed size, despite the decreased surface area to volume ratio. There was no relationship between avenacins and seed size except as measured in the elite panel in WI. Finally, relative concentration of AOSs (concentrated in the inner endosperm) decreased with seed size in the diversity panel but had no relationship to seed size in the elite panel. This relationship was further confirmed by examining the genetic correlation between seed traits and the specialized metabolites. In the diversity panel, there was strong positive genetic correlation between seed volume, seed surface area and hundred kernel weight and AVNs (>0.70), negative correlation with AOSs (< −0.23) and essentially no correlation with AECs (between 0 and −0.12). This relationship was less consistent when examined in the elite panel, and there were not consistent patterns between percent groat and metabolite traits in any panel or location (**Table S9**).

**Figure 2.**
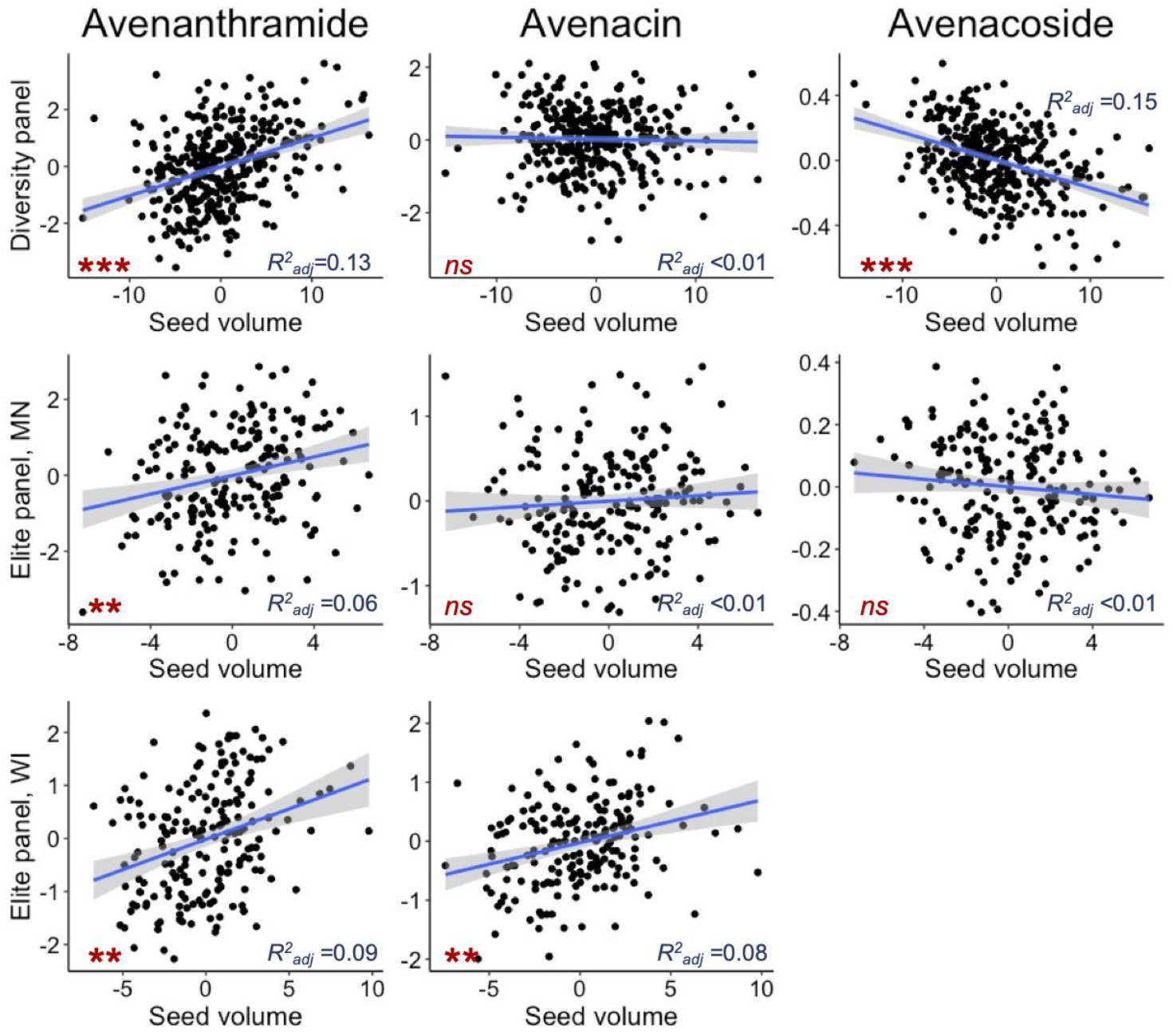
Relationship between specialized metabolites and seed size in the diversity panel (evaluated only in New York) and elite panel evaluated in Minnesota (“MN”) and Wisconsin (“WI”); data not available for the elite panel evaluated in South Dakota. For each metabolite class, an example was chosen where ‘Avenanthramide’ refers to avenanthramide B, ‘Avenacin’ refers to avenacin A1.1, and ‘Avenacoside’ refers to avenacoside A (**Table S1**). Model results for all metabolites are in presented **Table S7**. The *** indicates *p*<1e-6, ** *p*<1e-3, and ‘*ns*’ indicates *p*>0.05.

### Effect of breeding intensity on metabolites and seed traits

Using year of variety release as a proxy for plant breeding intensity (where later years indicate more intensive breeding efforts), we tested if breeding intensity affected seed size or metabolites in the individuals in the diversity panel for which these data are available (phenotypes and year information is available for 138 to 146 individuals per trait; **Table S10**). Seed volume increased over time and, correspondingly, seed surface area increased and the surface area to volume ratio decreased (**Figure 3a-c**). Hundred kernel weight and hundred hull weight both also increased over time, but groat percent remained constant (**Figure 3d-f**). Of the specialized metabolites, the relative concentration of avenacosides decreased over time, but avenanthramides and avenacins were unaffected (**Figure 3g-i**). Using multiple regression with year and seed volume as predictors for groat percentage and the specialized metabolites, the regression coefficient for year was not significantly different from zero for any metabolite (**Table S11**). These results indicate that while seed size was likely a target of selection as a yield component that had indirect effects on the seed metabolome composition, factors independent of size and breeding intensity also contributed to the observed metabolome variation.

**Figure 3.**
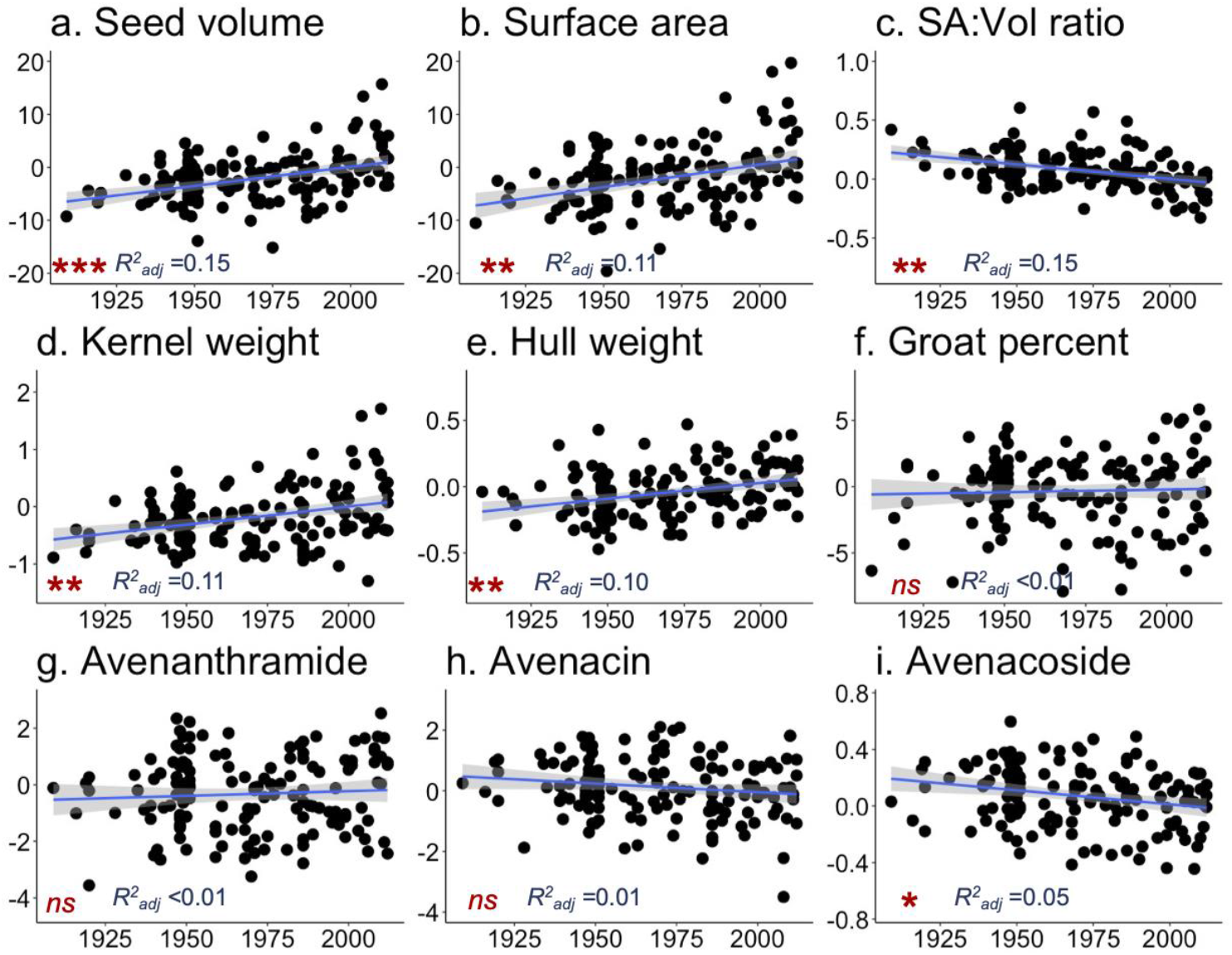
Relationship between year of variety release and deregressed BLUPs of (a) seed volume, (b) seed surface area, (c) seed surface area to volume ratio, (d) hundred kernel weight, (e) hundred hull weight, (f) groat percent, (g) avenanthramide, (h) an avenacin, and (i) an avenacoside in the diversity panel. For each metabolite class, an example was chosen where ‘Avenanthramide’ refers to avenanthramide B, ‘Avenacin’ refers to avenacin A1.1, and ‘Avenacoside’ refers to avenacoside A (**Table S1**). Model results for all traits are in presented **Table S10**. The *** indicates *p*<1e-6, ** *p*<1e-3, * *p*<0.05, and ‘*ns*’ indicates *p*>0.05.

### Genome wide association study

Single-trait GWAS was conducted for each of the specialized metabolites and seed traits in the diversity panel and each environment of the elite panel. Few metabolite traits had SNPs above a significance threshold of *p_Bonferroni_* < 0.05 (**Table 1**; **Figure S3**). No seed size traits had a significant GWAS result, but percent groat did in one environment of the elite panel (**Table 1**; **Figure S3**). None of these eleven significant SNPs were within genes (all genes within +/- 100kb of the SNPs are presented in **Table S12**). The significant GWAS results for AVN_A in the diversity panel on Chromosome 3A did not colocalize with known QTL for resistance to crown rust (McNish *et al*., 2020) (**Table S2**), despite the previously reported relationships between AVN concentration and crown rust resistance.

**Table 1.** Significant SNPs from GWAS of metabolites and seed traits by panel and environment. The diversity panel was evaluated in only one environment (NY, USA). The *P*-value is adjusted with a Bonferroni correction.

| Trait<sup>1</sup>, panel, environment | SNP | Chr, Position | *p*-value | Effect |
| --- | --- | --- | --- | --- |
| AVN_A, diversity, NY | avgbs_cluster_30159.1.28 | 3A, 406909563 | 0.035 | 0.34 |
| AEC_A1.1, diversity, NY | avgbs_32431.1.14 | 5A, 456500997 | 0.031 | -0.36 |
| AEC_A1.2, elite, SD | avgbs_cluster_3322.1.38 | 6C, 2212093 | 0.005 | -0.20 |
| AEC_A1.1, elite, WI | avgbs_21467.1.45 | 5D, 387376916 | 0.033 | 0.29 |
| AOS_dA, diversity, NY | avgbs_1891.1.28 | 4D, 266095186 | 0.038 | -0.25 |
| HKW, diversity, NY | avgbs_cluster_39333.1.13 | 2D, 518487763 | 0.002 | -0.41 |
| GP, elite, SD | avgbs_cluster_42433.1.28 | 3C, 3654644 | 0.007 | -3.24 |
| GP, elite, SD | avgbs_96083.1.13 | 3C, 3657557 | 0.015 | 3.36 |
| GP, elite, SD | avgbs_221727.1.25 | 3C, 6201470 | 0.008 | 3.23 |
| GP, elite, SD | avgbs_cluster_11404.1.64 | 3C, 7293210 | 0.013 | -3.18 |
| GP, elite SD | avgbs_cluster_11404.1.57 | 3C, 7293217 | 0.015 | -2.99 |

<sup>1</sup>Trait names are defined as follows: avenanthramide A , “AVN\_A”; avenanthramide B , “AVN\_B”; avenacin A1, “AEC\_A1.1” and “AEC\_A1.2”; avenacoside A, “AOS\_A”; 26-Desglucoavenacoside A, “AOS\_dA”; avenacoside B, “AOS\_B”

To visualize genomic regions relevant for metabolite and seed traits and determine if there is shared genetic control between traits, populations or environments, we examined all SNPs that met a reduced significance threshold of *p_FDR_* < 0.20 and plotted them in 10Mb bins (**Figure 4**). Within population and environment (e.g., elite panel in Minnesota), there were no shared SNPs between any two or more traits (e.g., between AVNs and seed size), indicating that the metabolite and seed traits do not have common large effect loci. Within AVNs, only results from the diversity panel met this threshold (**Figure 4a**). There were multiple points of overlap between environments and panels for AECs, with the highest count of shared SNPs on 5A (elite-MN, diversity panel) and 5C (elite-MN, elite-WI, diversity panel) (**Figure 4b**), and there were consistent SNPs identified for AOSs between the elite panel evaluated in MN and SD on chromosomes 1C and 4A (**Figure 4c**). However, the regions identified for seed size traits in the elite panel and the diversity panel were not shared (**Figure 4d**). As different genomic regions were implicated between panels and environments for the same trait, these results indicate genetic heterogeneity between panels and genotype-by-environment interactions.

**Figure 4.**
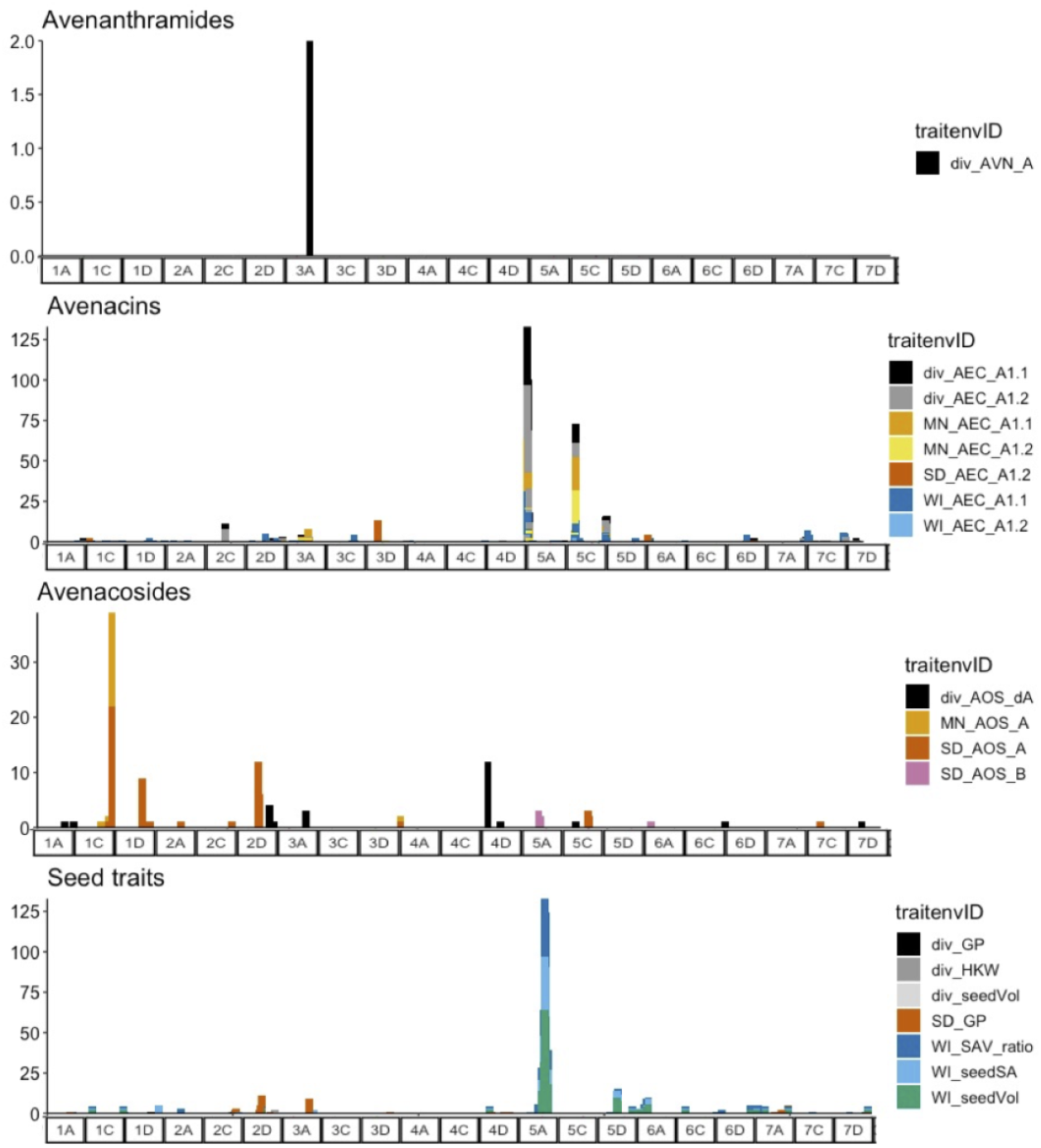
Number of SNPs from within 10Mb bins meeting a *p_FDR_* < 0.20 significance threshold from GWAS analysis by germplasm panel and environment. The panels show specific trait types (avenanthramides, avenacins, avenacosides and seed traits) where color indicates environment and specific trait.

### Transcriptome analyses

A transcriptome-wide association analysis (TWAS) was conducted for each of the specialized metabolites in the diversity panel to assess the relationship between gene expression and metabolite relative concentration. Of these, both AVNs had significant (*p_FDR_* < 0.05) TWAS results (72 for each AVN_A and AVN_B), with 51 shared and expression of most of these shared transcripts (50) positively correlated with increased AVNs (**Table 2, Table S13**). Of these, phenylalanine ammonia-lyase (“PAL”, TRINITY_DN26560_c0_g2_i1), the first committed enzyme of phenylpropanoid biosynthesis and phosphoenolpyruvate/phosphate translocator 1 (TRINITY_DN1581_c0_g1_i3), an enzyme in the pentose-phosphate pathway, a pathway that precedes the shikimate pathway could be connected to biosynthesis. The other specialized metabolites had few significant TWAS results (**Table 3**): the two AECs shared four significant transcripts and only AOS_B had a significant result. No significant transcripts were detected for any seed traits, even at a less stringent cutoff (*p_FDR_*<0.25).

**Table 2.**
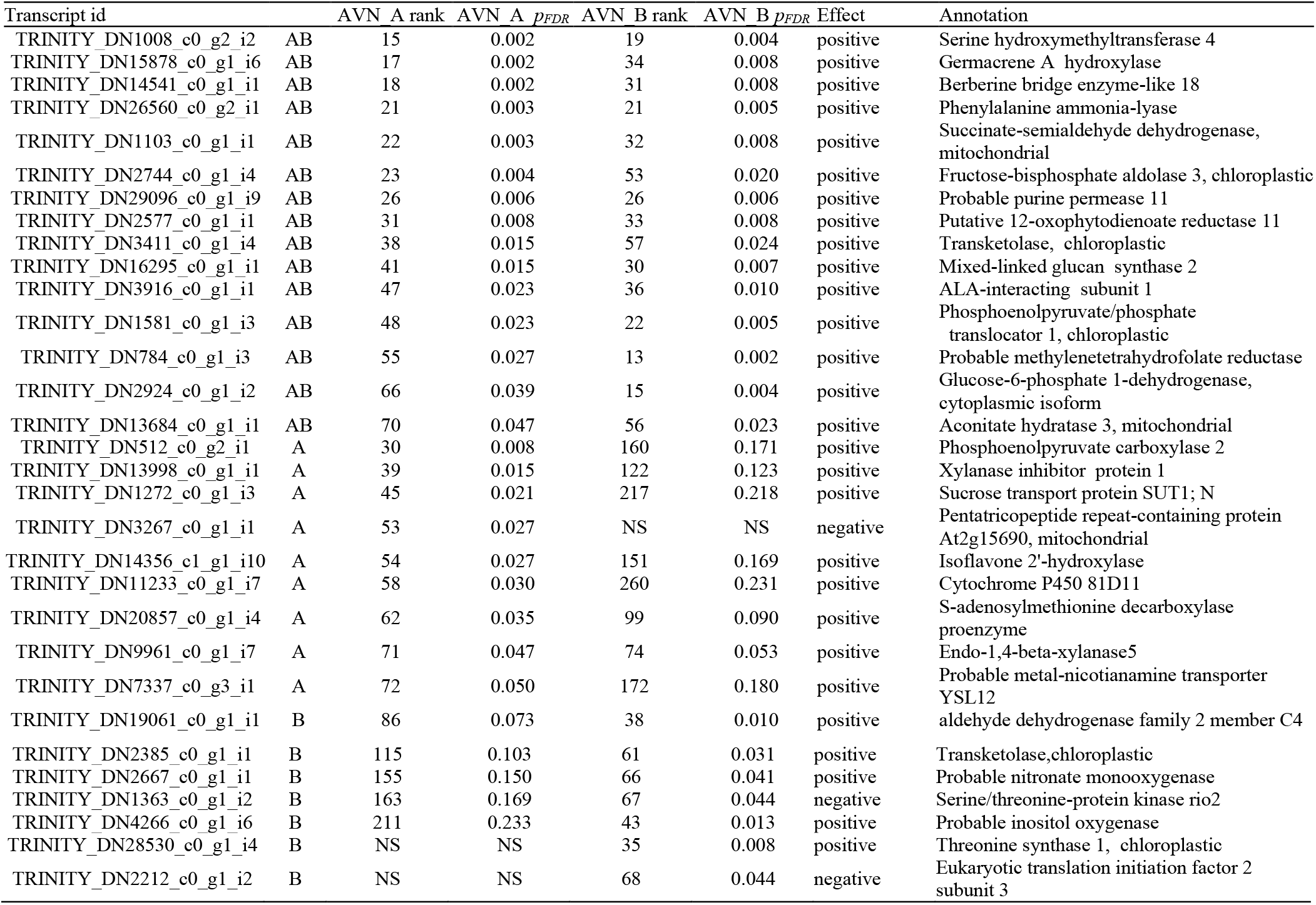
Significant transcripts (*p_FDR_* < 0.05) from TWAS of avenanthramides that have gene annotations. A full list of all significant transcripts is in **Table S13**. Rank refers to overall transcript significance in TWAS analysis, and effect refers to the direction of correlation between expression and relative concentration of avenanthramide.

| Transcript id | | AVN_A rank | AVN_A $p_{FDR}$ | AVN_B rank | AVN_B $p_{FDR}$ | Effect | Annotation |
| --- | --- | --- | --- | --- | --- | --- | --- |
| TRINITY_DN1008_c0_g2_i2 | AB | 15 | 0.002 | 19 | 0.004 | positive | Serine hydroxymethyltransferase 4 |
| TRINITY_DN15878_c0_g1_i6 | AB | 17 | 0.002 | 34 | 0.008 | positive | Germacrene A hydroxylase |
| TRINITY_DN14541_c0_g1_i1 | AB | 18 | 0.002 | 31 | 0.008 | positive | Berberine bridge enzyme-like 18 |
| TRINITY_DN26560_c0_g2_i1 | AB | 21 | 0.003 | 21 | 0.005 | positive | Phenylalanine ammonia-lyase |
| TRINITY_DN1103_c0_g1_i1 | AB | 22 | 0.003 | 32 | 0.008 | positive | Succinate-semialdehyde dehydrogenase, mitochondrial |
| TRINITY_DN2744_c0_g1_i4 | AB | 23 | 0.004 | 53 | 0.020 | positive | Fructose-bisphosphate aldolase 3, chloroplastic |
| TRINITY_DN29096_c0_g1_i9 | AB | 26 | 0.006 | 26 | 0.006 | positive | Probable purine permease 11 |
| TRINITY_DN2577_c0_g1_i1 | AB | 31 | 0.008 | 33 | 0.008 | positive | Putative 12-oxophytodienoate reductase 11 |
| TRINITY_DN3411_c0_g1_i4 | AB | 38 | 0.015 | 57 | 0.024 | positive | Transketolase, chloroplastic |
| TRINITY_DN16295_c0_g1_i1 | AB | 41 | 0.015 | 30 | 0.007 | positive | Mixed-linked glucan synthase 2 |
| TRINITY_DN3916_c0_g1_i1 | AB | 47 | 0.023 | 36 | 0.010 | positive | ALA-interacting subunit 1 |
| TRINITY_DN1581_c0_g1_i3 | AB | 48 | 0.023 | 22 | 0.005 | positive | Phosphoenolpyruvate/phosphate translocator 1, chloroplastic |
| TRINITY_DN784_c0_g1_i3 | AB | 55 | 0.027 | 13 | 0.002 | positive | Probable methylenetetrahydrofolate reductase |
| TRINITY_DN2924_c0_g1_i2 | AB | 66 | 0.039 | 15 | 0.004 | positive | Glucose-6-phosphate 1-dehydrogenase, cytoplasmic isoform |
| TRINITY_DN13684_c0_g1_i1 | AB | 70 | 0.047 | 56 | 0.023 | positive | Aconitate hydratase 3, mitochondrial |
| TRINITY_DN512_c0_g2_i1 | A | 30 | 0.008 | 160 | 0.171 | positive | Phosphoenolpyruvate carboxylase 2 |
| TRINITY_DN13998_c0_g1_i1 | A | 39 | 0.015 | 122 | 0.123 | positive | Xylanase inhibitor protein 1 |
| TRINITY_DN1272_c0_g1_i3 | A | 45 | 0.021 | 217 | 0.218 | positive | Sucrose transport protein SUT1; N |
| TRINITY_DN3267_c0_g1_i1 | A | 53 | 0.027 | NS | NS | negative | Pentatricopeptide repeat-containing protein At2g15690, mitochondrial |
| TRINITY_DN14356_c1_g1_i10 | A | 54 | 0.027 | 151 | 0.169 | positive | Isoflavone 2'-hydroxylase |
| TRINITY_DN11233_c0_g1_i7 | A | 58 | 0.030 | 260 | 0.231 | positive | Cytochrome P450 81D11 |
| TRINITY_DN20857_c0_g1_i4 | A | 62 | 0.035 | 99 | 0.090 | positive | S-adenosylmethionine decarboxylase proenzyme |
| TRINITY_DN9961_c0_g1_i7 | A | 71 | 0.047 | 74 | 0.053 | positive | Endo-1,4-beta-xylanase5 |
| TRINITY_DN7337_c0_g3_i1 | A | 72 | 0.050 | 172 | 0.180 | positive | Probable metal-nicotianamine transporter YSL12 |
| TRINITY_DN19061_c0_g1_i1 | B | 86 | 0.073 | 38 | 0.010 | positive | aldehyde dehydrogenase family 2 member C4 |
| TRINITY_DN2385_c0_g1_i1 | B | 115 | 0.103 | 61 | 0.031 | positive | Transketolase,chloroplastic |
| TRINITY_DN2667_c0_g1_i1 | B | 155 | 0.150 | 66 | 0.041 | positive | Probable nitronate monooxygenase |
| TRINITY_DN1363_c0_g1_i2 | B | 163 | 0.169 | 67 | 0.044 | negative | Serine/threonine-protein kinase rio2 |
| TRINITY_DN4266_c0_g1_i6 | B | 211 | 0.233 | 43 | 0.013 | positive | Probable inositol oxygenase |
| TRINITY_DN28530_c0_g1_i4 | B | NS | NS | 35 | 0.008 | positive | Threonine synthase 1, chloroplastic |
| TRINITY_DN2212_c0_g1_i2 | B | NS | NS | 68 | 0.044 | negative | Eukaryotic translation initiation factor 2 subunit 3 |

**Table 3.** Significant transcripts (*p_FDR_* < 0.05) from TWAS of avenacins (AEC) and avenacosides (AOS) where rank refers to overall transcript significance in TWAS analysis, and effect refers to the direction of correlation between expression and relative metabolite concentration. Annotations are provided when available.

| Transcript name | AEC_A1.1 |  | AEC_A1.2 |  | AOS_B |  | Direction | Annotation |
| --- | --- | --- | --- | --- | --- | --- | --- | --- |
| | rank | $p_{FDR}$ | rank | $p_{FDR}$ | rank | $p_{FDR}$ | | |
| TRINITY_DN36363_c0_g2_i1 | 1 | 4.1E-05 | 1 | 8.E-05 | - | - | positive |  |
| TRINITY_DN6771_c0_g1_i1 | 2 | 2.4E-04 | 2 | 0.006 | - | - | positive | Phosphoethanolamine N-methyltransferase 1 |
| TRINITY_DN97809_c0_g1_i1 | 3 | 0.03 | 4 | 0.04 | - | - | positive |  |
| TRINITY_DN7675_c0_g1_i7 | 5 | 0.09 | 3 | 0.02 | - | - | positive |  |
| TRINITY_DN1526_c0_g1_i12 | - | - | - | - | 1 | 0.050 | negative |  |

To better understand the biological relevance of the rest of the transcripts, GO enrichment analysis was conducted on the false-discovery rate adjusted *p*-values. While only AVN_B had a significantly enriched term after multiple test correction (pentose-phosphate shunt, GO:0006098), GO terms related to the shikimate (chorismate biosynthetic process, GO:0009423) and L-phenylalanine catabolic processes (GO:0006559) were top GO terms for all avenanthramides (**Table 4**). There was no significant enrichment of GO terms for either the avenacins (**Table S14**) or avenacosides (**Table S15**).

**Table 4.** GO enrichment of biological process terms for avenanthramide TWAS results where the top three GO terms from each avenanthramide (AVN) are presented along with the rank for the other avenanthramides. The p-values are unadjusted and the * indicates that it is significant when adjusted for a false discovery rate.

| GO ID | Term | AVN_A |  | AVN_B |  |
| --- | --- | --- | --- | --- | --- |
|  |  | rank | *p* | rank | *p* |
| GO:0006098 | pentose-phosphate shunt | 1 | 6.1E-05 | 1 | 7.7E-06* |
| GO:0006559 | L-phenylalanine catabolic process | 2 | 2.8E-04 | - | - |
| GO:0009423 | chorismate biosynthetic process | 3 | 7.1E-04 | 2 | 1.7E-03 |
| GO:0090630 | activation of GTPase activity | - | - | 3 | 2.5E-03 |

We also examined how expression of the significant TWAS transcripts changed over seed development for AVNs. Developing oat seed transcripts were categorized into temporally covarying groups (Hu *et al*., 2020) and we found that significant transcripts from AVN TWAS analysis were enriched for transcripts that had a trajectory of increased expression beginning at eight days after anthesis when compared to all transcripts (hypergeometric test, AVN_A: *p*=2.39e-13, AVN_B: *p*=2.82e-10). In contrast, there was weak evidence for enrichment of any transcript class in known avenanthramide biosynthetic enzymes (hypergeometric test, decrease in expression at 8DAA, *p*= 0.049) or Shikimate pathway enzymes (hypergeometric test, decrease in expression at 13DAA, *p*= 0.062) (**Figure 5**).

**Figure 5.**
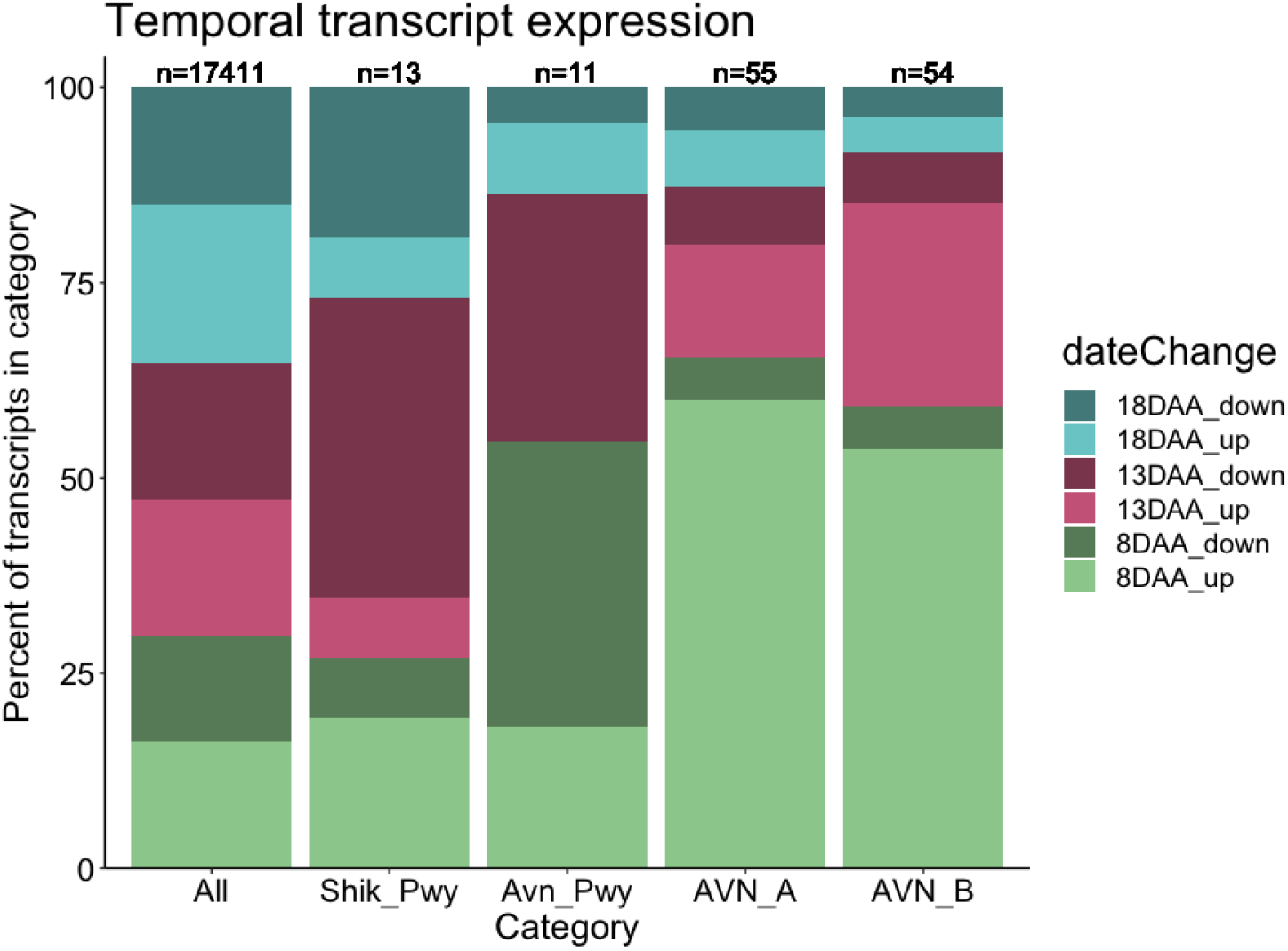
Oat seed transcripts classified by temporal variant category and direction as described in (Hu *et al*., 2020). The percent of transcripts in each category is shown for all transcripts in the dataset (“all”), transcripts annotated to be part of the preceding shikimate pathway (“Shik_Pwy”), transcripts annotated in avenanthramide biosynthesis (“Avn_Pwy”), and each of the avenanthramides (A, B and C). The numbers at the top indicate the number of transcripts that were annotated by temporal group.

Finally, we tested if seed volume corresponded to expression of AVN TWAS results to determine if there was expression variation independent of seed volume that could be a target of selection. Seed size was less predictive of TWAS gene expression than the phenotype (AVN_B) as measured by coefficient of determination (**Table S16**). For instance, PAL and Phosphoenolpyruvate/Phosphate Translocator 1 were not strongly associated with seed volume (**Figure 6)**. These results indicate that while relative concentration of AVN tracks with seed volume and gene expression, gene expression is not strongly linked to seed volume, and thus gene expression is an independent contributor to patterns of variation in AVN abundance.

**Figure 6.**
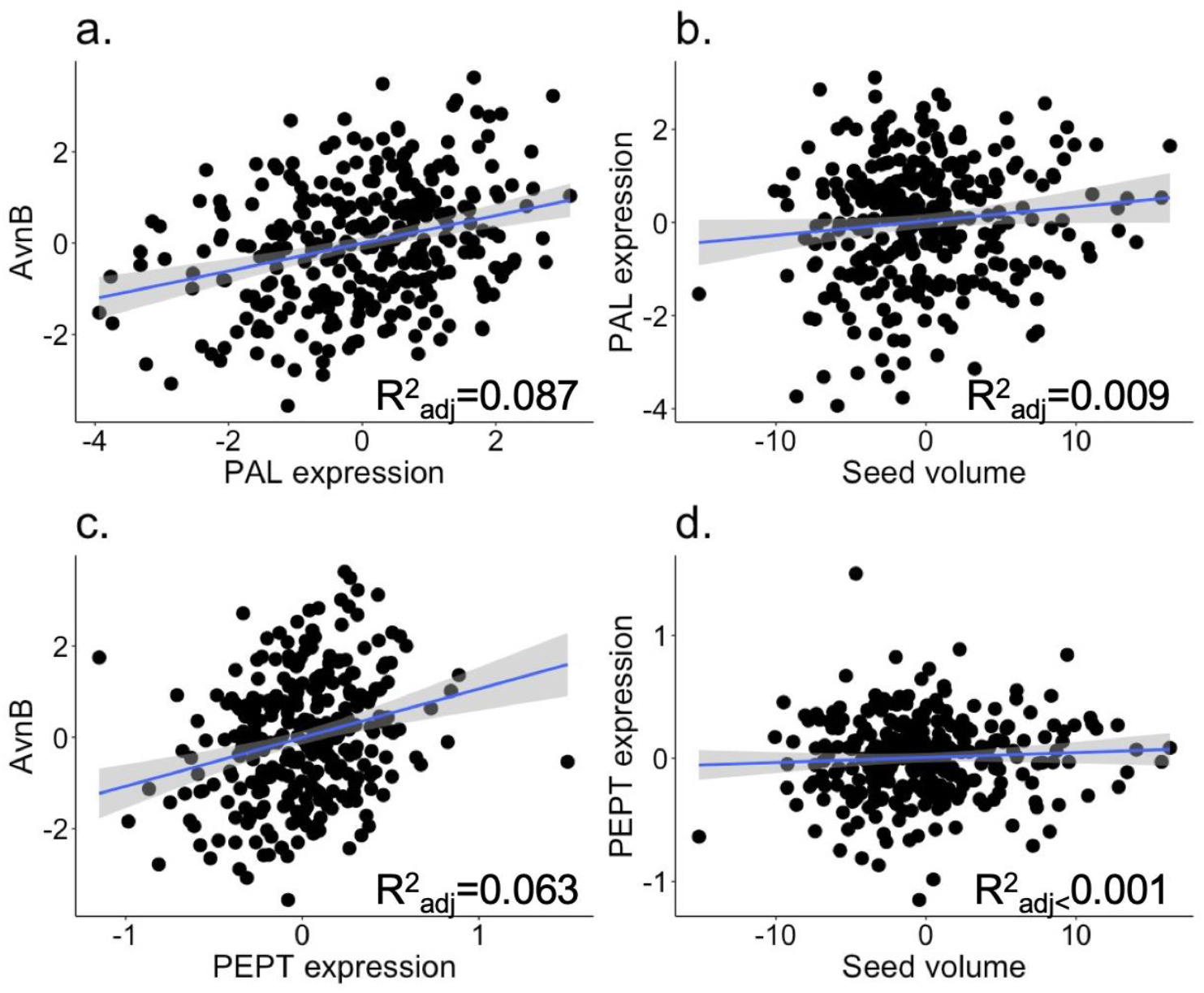
The relationship and coefficient of determination between expression of (a) phenylalanine ammonia-lyase, “PAL” and (c) phosphoenolpyruvate/phosphate translocator 1, “PEPT” and avenanthramide B (“Avn_B”) concentration and the relationship between seed volume and (b) PAL and (d) PEPT expression. The relationship between avenanthramide B and all TWAS results are given in **Table S16**.

### eQTL analysis

Because we predicted that expression variation is important for oat specialized metabolites, especially AVNs, we conducted eQTL analysis on genes detected in TWAS and on known pathway genes and examined if those eQTL colocalized with our GWAS results. Two avenanthramide TWAS results had eQTL at a *p_FDR_*<0.05 threshold, TRINITY_DN1008_c0_g2_i2 a serine hydroxymethyltransferase 4, and TRINITY_DN13684_c0_g1_i1 a mitochondrial aconitate hydratase 3. These two genes neither co-localized with the avenanthramide GWAS result nor were definitively annotated to a single position in the oat genome. Relaxing the significance threshold to *p_FDR_*<0.2 revealed eQTL of four additional genes (**Figure S4**), but the eQTLs detected on chromosome 3A were not in LD with the GWAS result (*r^2^* < 0.02 for all).

Of the pathway genes (**Table S5**), only TRINITY_DN2726_c0_g1_i2, a bifunctional 3-dehydroquinate dehydratase/shikimate dehydrogenase, had a significant eQTL (*p_FDR_* = 0.002; Chr 3A, position 15737366, avgbs_cluster_12707.1.49). We also examined eQTL from pathway genes at a *p_FDR_*<0.2 threshold and identified eQTL of five additional genes (**Figure S4**). We found that eQTL of TRINITY_DN1661_c0_g1_i1, an anthranilate synthase (avenanthramides are a condensation between phenolic acids and anthranilic acid), was in LD with the avenanthramide GWAS result on chromosome 3A with the strongest association being the SNP avgbs_cluster_34200.1.64 (*p_FDR_* = 0.16, *r^2^* = 0.44).

## Discussion

Oat (*Avena sativa* L.) is a cereal crop with known health benefits from consuming the grain or through topical skincare application. These benefits are derived from a diverse suite of metabolites, including unsaturated fatty acids and ß-glucans as well as the specialized avenanthramides, avencins and avenacosides. We characterized the genomic and transcriptomic bases of specialized metabolite variation in diverse and elite oat germplasm in the context of seed size and selection over a century of oat breeding. We found that variation is diminished in elite germplasm, but selection for larger seeds only accounts for part of that reduction. For avenanthramides in particular, we found in addition to increased abundance in larger seeds, there was also variation in biosynthetic enzymes upstream of the committed pathway enzymes that contributed to phenotypic variation. Broadly, this work addresses longstanding questions about how crop breeding has shaped specialized metabolome profiles, and prospects for continued plant breeding.

### Historical dimensions of oat specialized metabolism and change in seed size

Specialized metabolites serve multiple purposes in plants, with one prominent use being plant defense against biotic stresses (Mithöfer & Boland, 2012; Kessler & Kalske, 2018; Jacobowitz & Weng, 2020). The relationship between plant domestication and breeding, resistance to biotic stress, and specialized metabolites has been widely examined to understand how plant selection has shaped agro-ecological interactions. Most work has been conducted comparing wild and domesticated plants, and has found that cultivated plants are more susceptible to biotic stress than their wild progenitors (Turcotte *et al*., 2014; Whitehead *et al*., 2017; Fernandez *et al*., 2021). A concomitant decrease in secondary metabolites, however, has not been consistently observed (Whitehead *et al*., 2017). Instead, tradeoffs between plant growth and defense (Whitehead & Poveda, 2019) or plant nutrition (Fernandez *et al*., 2021) may be important factors. The studies that interrogate a spectrum of plant breeding intensity from domestication to landraces to modern varieties have used less than 25 accessions each, and have produced mixed results where some find a decrease in resistance with breeding intensity (Rosenthal & Dirzo, 1997; Lindig-Cisneros *et al*., 2002) but others do not (Ferrero *et al*., 2020). Intriguingly, Lindig-Cisneros *et al*., (2002) associated reduced biotic stress resistance with reduced metabolite diversity, but not absolute metabolite concentrations. Overall, these findings indicate that there are nuanced crop-specific patterns in how breeding has shaped specialized metabolites (and plant defense), but there is a need for work that includes a greater number of plant accessions and a finer-scale gradient of plant breeding intensity.

In our work, we surveyed oats spanning almost a century of plant breeding - beginning with the rediscovery of Mendel in the early 20th century to genomics-enabled breeding in the 21st century. Yield has consistently been a trait of plant breeding interest, with yield gains throughout the 20th century (Rodgers *et al*., 1983) and is still a focus of current breeding programs (Haikka *et al*., 2020; González-Barrios *et al*., 2021).We examined the relationship between breeding intensity (by year of variety release), seed size, and defensive metabolites in more than 138 individuals. We found that more intensive breeding led to larger oat seeds, but not a greater proportion of edible tissue (groat) and, while relative concentrations of specialized metabolites were tied to seed size, they were not a direct target of plant breeding. We found that larger seeds had high avenanthramide abundance, despite decreased surface area to volume ratio inherent to larger seeds, but there was no relationship with breeding intensity. In contrast, avenacoside abundance decreased with increasing seed size associated with breeding intensity, despite larger endosperm volume. These results indicate that there are not consistent tradeoffs between growth (seed size) and defense (avenanthramides, avenacosides). Further, we found that ongoing plant breeding did not uniformly reduce or increase plant specialized metabolites, but may have affected size of and concentration of metabolites in specific seed tissues (like the aleurone layer).

### Breeding for oat avenanthramides

Of the oat seed specialized metabolites, avenanthramides have garnered the most research interest. Avenanthramides are antioxidants (Bratt *et al*., 2003) and have been implicated in resistance to the oat crown rust (Wise *et al*., 2008; Wise, 2014). The avenanthramide biosynthetic pathway has been defined (Collins, 2011; Wise, 2014; Li *et al*., 2019), yet this work has not been translated into tools for oat breeders, like molecular markers. Critically, it remains unknown whether functional or regulatory mutations in the committed biosynthetic pathway enzymes (enzymes specific to avenanthramide biosynthesis) or upstream biosynthetic pathway enzymes (not specific to avenanthramide biosynthesis) are the most significant contributors to heritable variation in cultivated oat germplasm. Neither our GWAS nor TWAS results implicated committed pathway genes. Instead, TWAS revealed that biosynthetically upstream enzymes expressed early in seed development contributed to avenanthramide abundance. In addition, we found that an eQTL of a biosynthetically upstream enzyme co-localized with our avenanthramide GWAS result. While our interpretation and enrichment analyses were limited by availability of transcript annotations (which, likely, are more complete for highly conserved, rather than oat-specific, genes) these results nonetheless suggest that regulation of or flux through the pathway may be a promising avenue for plant breeding.

Dimberg & Peterson (2009) examined the relationship between avenanthramides and compounds that are precursors or derived from other branches of related biosynthetic pathways. Their results did not offer a straightforward indication of which biosynthetic step moderates pathway flux; instead, PAL expression did not depend upon the amount of its substrate (phenylalanine) nor did PAL expression affect expression of HHT (the terminal enzyme in avenanthramide biosynthesis). Our results implicate PAL expression as important for avenanthramide abundance, as well as a phosphoenolpyruvate translocator in the pentose phosphate pathway, and other transcripts of unknown function. These results add to the widely recognized importance of PAL expression as a regulator of flux in phenylpropanoid biosynthesis (Huang *et al*., 2010; Kim & Hwang, 2014; Barros & Dixon, 2020). In addition, a broader examination of precursor metabolites, including those in the pentose phosphate pathway may produce interesting results as diversification of enzymes from primary metabolism is important for contributing to specialized metabolism diversity (Moghe & Last, 2015; Maeda, 2019). Overall, our results should prompt future work on avenanthramides to focus on upstream biosynthetic processes, as most variation affecting avenanthramides appears to be in enzymes preceding committed biosynthetic steps.

Our results also contribute to an understanding of when avenanthramide biosynthesis occurs in oat seeds. Avenanthramides are detected as early as eight days after anthesis (DAA), and while Hu *et al*. (2020) found that HHT is expressed at 8 DAA, Peterson & Dimberg, (2008) did not observe expression until 20 DAA. By sampling gene expression at only 23 DAA, we likely sampled at a time where it would be possible to detect differences in HHT expression, but we may have missed peak differential expression of upstream enzymes that contributed pathway flux. Our avenanthramide TWAS results were enriched for genes that were expressed early in seed development (8 DAA), and Hu *et al*., (2020) found that two other pathway enzymes, 4-coumaroyl-CoA3-hydroxylase (CCoA3H), caffeoyl-CoA3-O-methyltransferase (CCoAOMT) increase in expression early in development before dropping beginning at 18 DAA. Together, these results indicate that the precursors of avenanthramides may be biosynthesized early in seed development. Our understanding will improve with further use of oat genomic resources, as well as transcriptomic analysis paired with metabolomic profiling over seed development.

Finally, despite the connection between avenanthramides and the disease, crown rust, no results from our GWAS or TWAS results colocalized with previously reported crown rust QTL (McNish *et al*., 2020). One explanation for this finding is that we did not inoculate oats with crown rust, nor trigger systemic acquired resistance (SAR). Both crown rust infection and treating oats with analogs of hormones that activate SAR increase avenanthramide concentration (Wise *et al*., 2008, 2016; Wise, 2011, 2017). We predict that, if SAR was activated, there would be more extreme variation in avenanthramide concentrations and we would implicate more genetic loci, some of which would colocalize with crown rust QTL due to shared regulation. Overall, these results suggest that genetic variation in regulation exists, but regulatory elements may need to be activated to effectively map or select upon this variation.

### Prospects for oat saponins – avenacins and avenacosides

The saponins of oats are of interest from a human health perspective as they are associated with reduction of cholesterol (Sang & Chu, 2017). Our results did not implicate promising candidate genes by GWAS nor TWAS that could be applied to develop tools for plant breeders. Like avenanthramides, our TWAS results are limited by only sampling at one time point. We also found that the saponins, especially the avenacosides, were more sporadically detected in the elite germplasm and within compound class correlations were weaker, potentially indicating a decrease in abundance in moving from diverse to elite germplasm. This may be due to taste: high concentrations of avenacosides in oat seed can contribute to an undesirable bitter off taste (Günther-Jordanland *et al*., 2016, 2020). Selection for organoleptic quality has been implicated in reducing saponin concentration in cultivated legumes (Ku *et al*., 2020), and our results indicate there has been a similar historical trajectory in oat. However, to the best of our knowledge, current oat breeding efforts do not regularly incorporate sensory evaluations.

### Selection for an optimized oat seed specialized metabolome

In breeding for nutrition, flavor, or aesthetics (color), plant breeders have changed crop metabolomic profiles. However, working with specialized metabolites versus major nutritional metabolites presents different challenges and thus may require different plant breeding approaches. As an example, fatty acid methyl esters (FAMEs) are healthful fats in oat seed that comprise 3-11% of oat seed composition, compared to 0.2% for avenanthramides. Also, while fatty acid biosynthetic enzymes have some degree of cross-species conservation, this is not true for avenanthramides that are only present in a few (non-model) plant species (Ponchet *et al*., 1988; Wise, 2014) and a caterpillar (Blaakmeer *et al*., 1994). In addition, the specialized metabolites we measured in oats are negatively correlated and do not have shared genetic control, presenting a challenge for selecting for both traits simultaneously but promising for efforts to select for a single trait. Finally, and perhaps most importantly, the specialized metabolite heritability (AVNs: h^2^ < 0.26, AECs: h^2^ < 0.61, AOSs: h^2^ < 0.52) we report here is lower than that of FAMEs (h^2^ > 0.61) (Carlson *et al*., 2019). Overall, these results suggest that work to increase specialized metabolite concentrations will benefit from strategies that reduce environmental variation to improve trait heritability, or increase replication in plant breeding trials, and incorporate seed size into phenotyping efforts.

### Conclusions

An understanding of patterns of variation in the plant specialized metabolome is relevant for developing health-promoting functional food crops that may also withstand biotic stress. Due to the low concentrations and lineage specificity of specialized metabolites, they are infrequent direct targets of plant breeding, but may have been inadvertently shaped through processes like selection on other traits or genetic drift. In a diverse panel of cultivated oats, we measured seed size and specialized metabolites and conducted genomic and transcriptomic analyses to characterize existing variation and the processes that contributed to it. Overall, we show that the increased seed size associated with modern plant breeding has uneven effects on the oat seed metabolome, and variation also exists independently of seed size. Broadly, despite the multitudes of phenotypic changes in crops from plant breeding, variation for some specialized metabolites persists in cultivated plants and could be targeted by future plant breeding efforts.

## Acknowledgements

Funding for this research was provided by USDA-NIFA-AFRI 2017-67007-26502 and Hatch Project 149-577.

## Author Contributions

JLJ, MAG and MES designed the research. LJB, HH, and MTC analyzed the data, and HH, MTC, MCT, LG, KPS and MES conducted experiments. CB conducted the metabolomics extraction, measurement, and data processing. LJB, MAG and JLJ wrote the manuscript and all co-authors were involved in editing the manuscript.

## Data Availability

Data files used in these analyses, including all metabolite and seed trait phenotypes for all panels and locations, gene expression data, gene annotations and genotypes are publicly available in CyVerse(https://datacommons.cyverse.org/browse/iplant/home/shared/GoreLab/dataFromPubs/Brzozowski_OatMetabolome_2021). Associated raw data has been previously published to public repositories (Metabolomics: (Haikka *et al*., 2020); gene expression:https://datacommons.cyverse.org/browse/iplant/home/shared/commons_repo/curated/HaixiaoHu_OatMultOmicsPred_Jun2021 (DOI: 10.25739/8p1e-0931). Scripts used in this work are available in github (https://github.com/ljbrzozowski/OatSeed_SpecializedMetabolomics).

## Supporting information contents

**Figure S1.** Scree plots for genomic principal component analysis

**Figure S2**. Scree plots for PEER analysis of gene expression data

**Figure S3.** Genome-wide association study results for all metabolites, and germplasm panels with significant results

**Figure S4.** eQTL analysis results from genes implicated in avenanthramide TWAS analysis and known biosynthetic genes

**Table S1.** Metabolite spectra and names in both germplasm panels

**Table S2**. ANOVA results for days to heading (DTH) covariate significance in drBLUP calculation for metabolite and seed size phenotypes

**Table S3**. Year of variety release from all available oat lines in diversity panel

**Table S4**. Crown rust QTL SNPs mapped to most recent genome

**Table S5**. Transcripts associated with avenanthramide biosynthetic pathway

**Table S6**. Seed size heritability

**Table S7**. Relationship between seed size and relative metabolite concentration ANOVA results

**Table S8.** Relationship between seed weight and metabolite relative metabolite concentration ANOVA results

**Table S9**. Genetic correlation between seed size and specialized metabolites

**Table S10.** Relationships between variety release year and seed size and metabolite relative metabolite concentration ANOVA results.

**Table S11**. Results from multiple regression analysis using variety release year and seed size.

**Table S12**. All genes within 100kB of significant GWAS results

**Table S13**. Full avenanthramide transcriptome-wide association study (TWAS) results

**Table S14**. GO enrichment of biological process terms from avenacin TWAS results

**Table S15**. GO enrichment of biological process terms from avenacoside TWAS results

**Table S16**. ANOVA results for relationship between gene expression and seed volume

**Method S1**. Metabolite extraction, measurement and annotation

