## Supporting Information for "Selection for seed size has indirectly shaped specialized metabolite abundance in oat (*Avena sativa* L.)"

The following Supporting Information is available for this article:

**Fig. S1** Scree plots for genomic principal component analysis

**Fig. S2** Scree plots for PEER analysis of gene expression data

**Fig. S3** Genome-wide association study results for all metabolites, and germplasm panels with significant results

**Fig. S4** eQTL analysis results from genes implicated in avenanthramide TWAS analysis and known biosynthetic genes

**Table S1** Metabolite spectra and names in both germplasm panels

**Table S2** ANOVA results for days to heading (DTH) covariate significance in drBLUP calculation for metabolite and seed size phenotypes

**Table S3** Year of variety release from all available oat lines in diversity panel

**Table S4** Crown rust QTL SNPs mapped to most recent genome

**Table S5** Transcripts associated with avenanthramide biosynthetic pathway

**Table S6** Seed size heritability

**Table S7** Relationship between seed size and relative metabolite concentration ANOVA results

**Table S8** Relationship between seed weight and metabolite relative metabolite concentration ANOVA results

**Table S9** Genetic correlation between seed size and specialized metabolites

**Table S10** Relationships between variety release year and seed size and metabolite relative metabolite concentration ANOVA results

**Table S11** Results from multiple regression analysis using variety release year and seed size

**Table S12** All genes within 100kB of significant GWAS results

**Table S13** Full avenanthramide transcriptome-wide association study (TWAS) results

**Table S14** GO enrichment of biological process terms from avenacin TWAS results

**Table S15** GO enrichment of biological process terms from avenacoside TWAS results

**Table S16** ANOVA results for relationship between gene expression and seed volume

**Methods S1** Metabolite extraction, measurement and annotation

**Fig. S1** Scree plots for principal component analysis of GBS data.

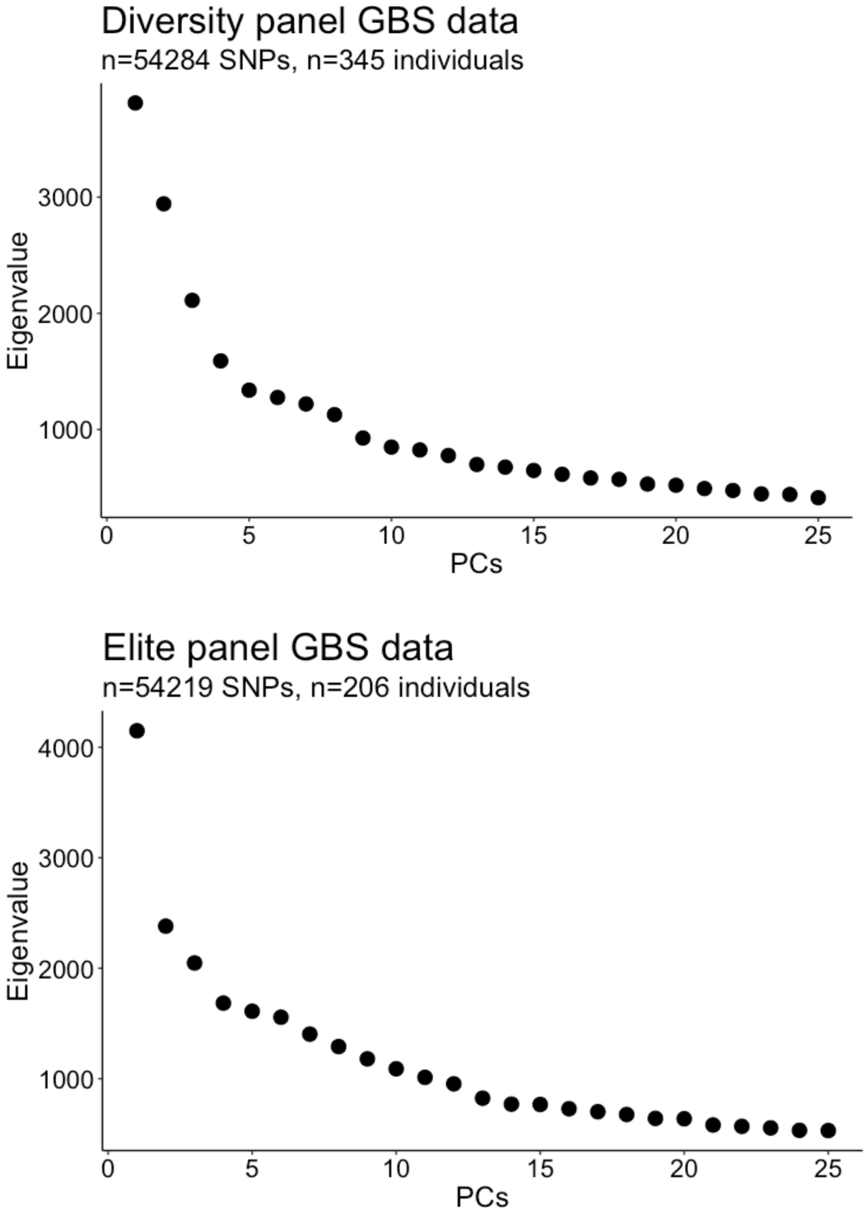

**Fig. S2** Scree plots for PEER analysis of gene expression data.

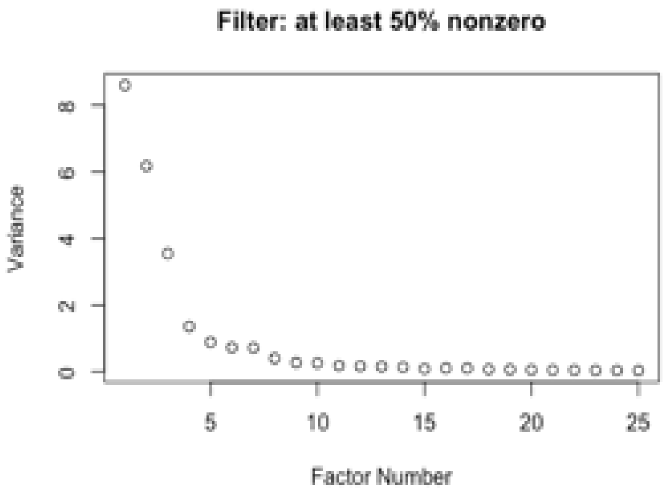

**Fig. S3** Genome-wide association study results (Manhattan plots and QQ-plots) for all metabolites, and germplasm panels with significant results (**Table 1**).

a. Avenanthramide A (LC.03.1204), Diversity panel

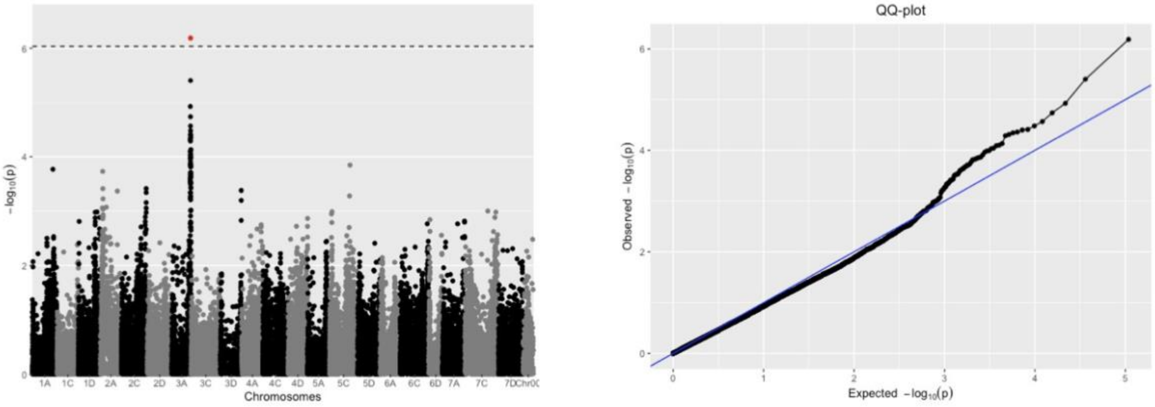

b. Avenacin A1.1 (LC.03.0066), Diversity panel

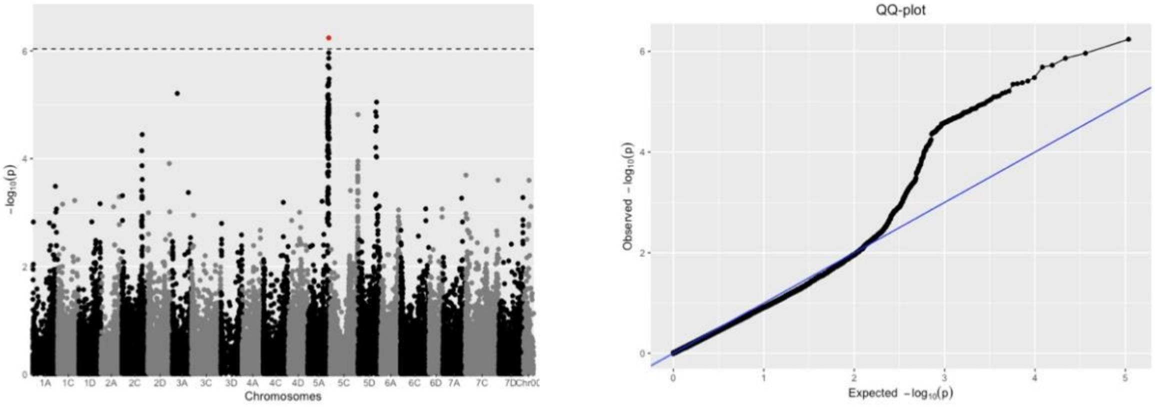

c. 26-Desglucoavenacoside A (LC.03.0052), Diversity panel

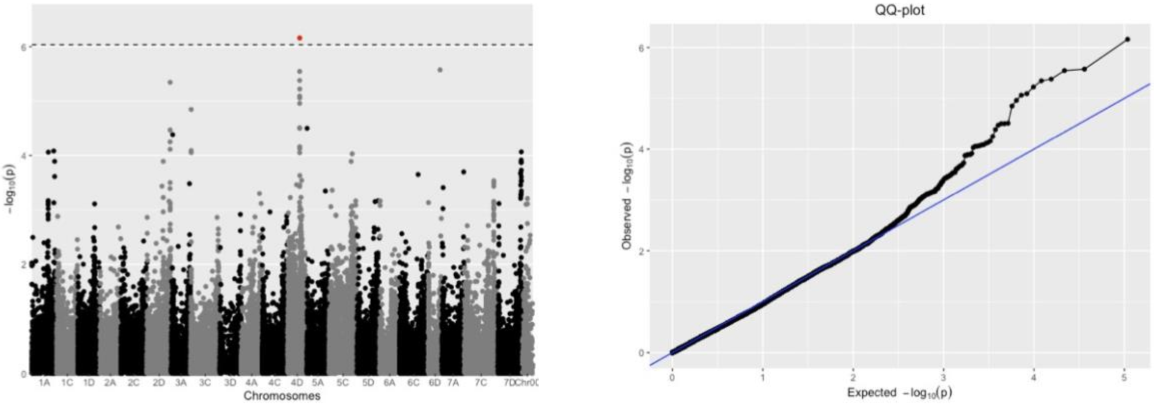

d. Avenacin A1.2 (LC.02.0051), Elite panel, SD

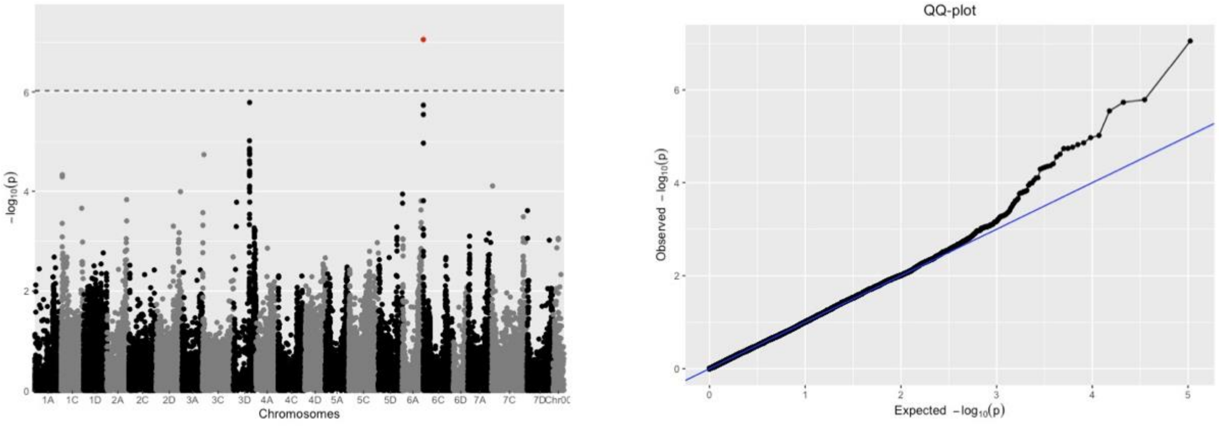

e. Avenacin A1.1 (LC.02.0099), Elite panel, WI

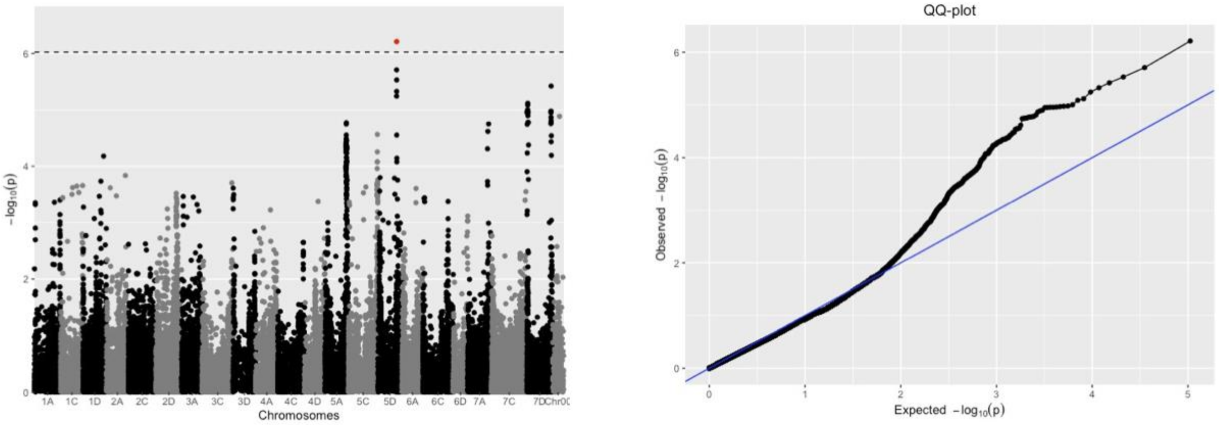

f. Hundred kernel weight (HKW), Diversity panel

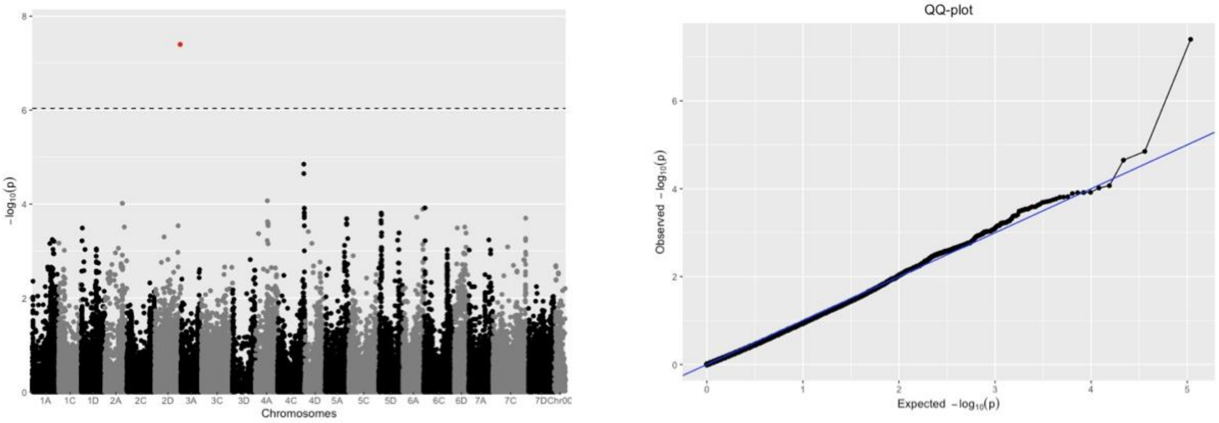

g. Groat percent (GP), Elite panel, SD

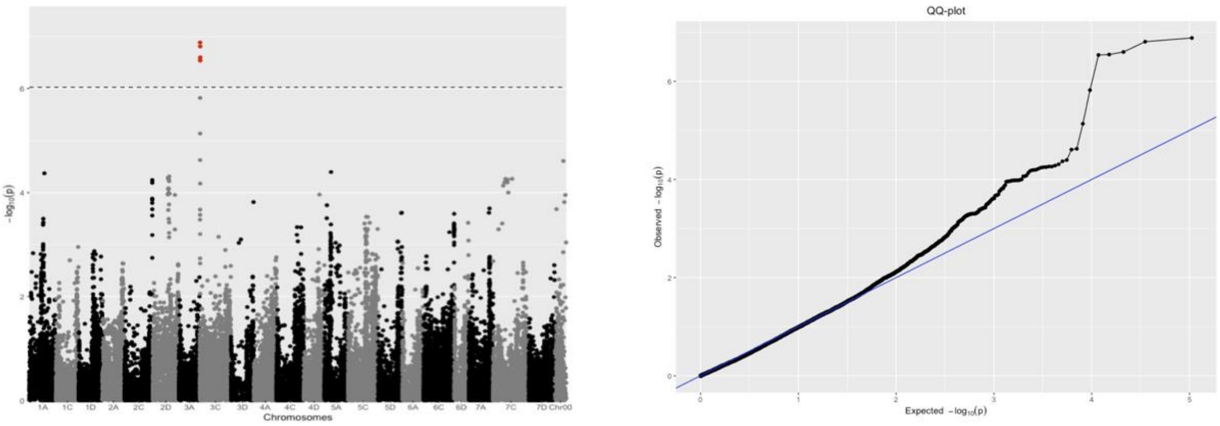

**Fig. S4** eQTL results from the genes implicated in (a) avenanthramide TWAS analysis ($p_{FDR}$<0.05 for both AVN_A and AVN_B) and those hypothesized to be in the biosynthetic pathway (Table S5). Each point is an eQTL, and only eQTL with $p_{FDR}$<0.20 are shown.

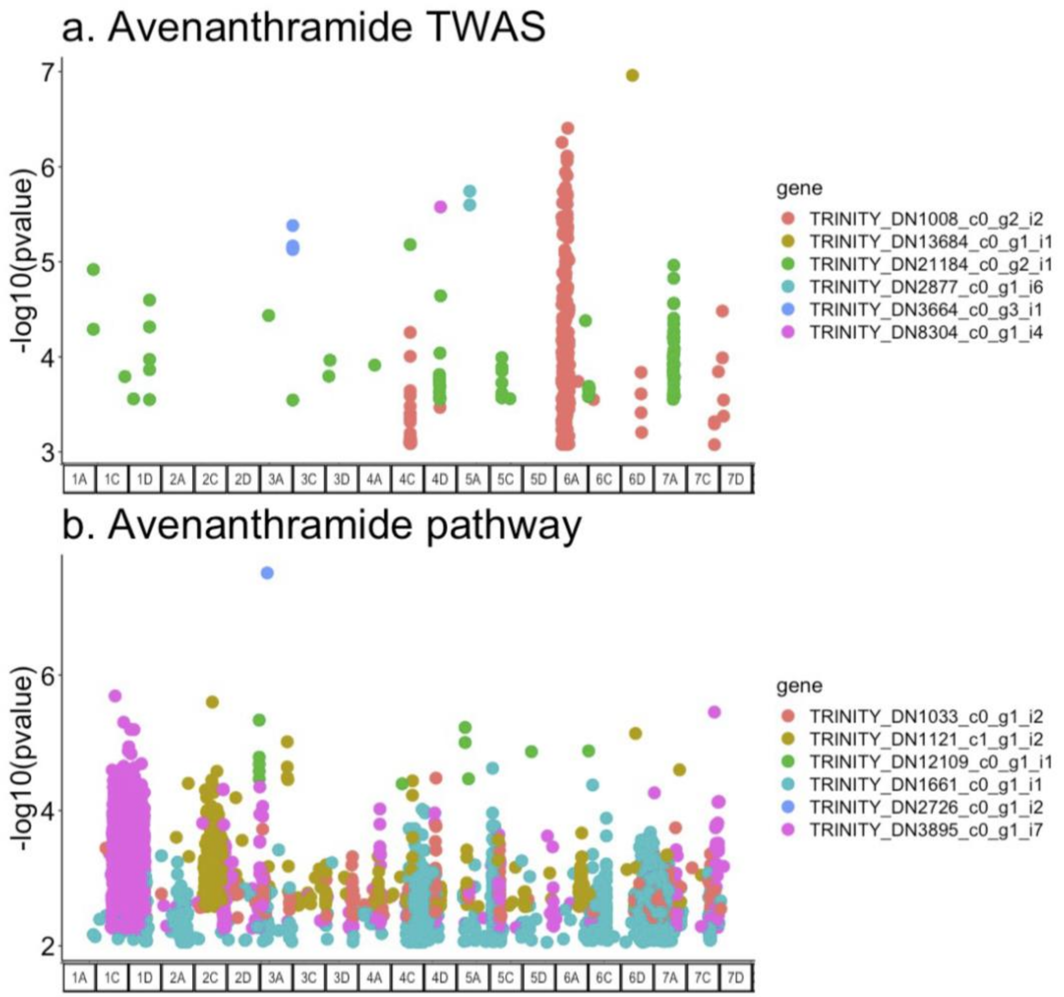

**Table S1** All specialized metabolites (two avenanthramides, two avenacins, and three avenacosides) were annotated by RamClustR in the diversity panel. In the elite panel, measured previously separately from the diversity panel, all avenanthramides and one avenacin was annotated. As some individuals were measured with both panels, the specialized metabolites detected in the diversity panel should also be present in the elite panel. To identify these metabolites in the elite panel, we matched mass to charge ratio (*m/z*) where shared *m/z* are indicated by bolded numbers. For compounds with more than 12 peaks, the 12 most prominent (largest absorbance) are noted. An asterisk (*) indicates that these characteristic *m/z* are also described in the literature. For the avenanthramides, characteristic *m/z* from positive mode MS2 are given in Table S2 of de Bruijn *et al.*, (2019). For the avenacins, a major *m/z* peak is the mass of an acyl group greater than the unacylated avenacins in Figure S1 in Leveau *et al.*, (2019). Characteristic *m/z* peaks of the avenacosides are given in Bahraminejad *et al.*, (2008).

| Category | Annotation | Diversity panel | | Elite panel | |
|---|---|---|---|---|---|
| | | Annotated? | *m/z* | Annotated? | *m/z* |
| Avenanthramides | Avenanthramide A (AVN_A) | yes, LC.03.1204 | **300**, 355 | yes, LC.02.0325 | 147*, 148, **300**, 322, 119, 91 |
| | Avenanthramide B (AVN_B) | yes, LC.03.0125 | 657, 504, **330**, 288, 658, 505, 639, 353, 323, 179, 385, 362 | yes, LC.02.0242 | 117, 145, 177*, 89, 149, 352, 178, **330** |

| | | | | | |
|---|---|---|---|---|---|
| Avenacosides | Avenacoside A (AOS_A) | yes, LC.03.0001 | **431***, **413***, **271**, **432**, **593***, 253, 414, 755, **739***, **594**, **309**, **272** | no, LC.02.0012 | **431***, **413***, **271**, **432**, **593***, 414, **309**, **594**, **739***, **272**, 433, 129, |
| | 26-Desglucoavenacoside A (AOS_dA) | yes, LC.03.0052 | **739**, **923**, **721**, **740**, **924**, **722**, **741**, 271, 579, **925**, 723, **901*** | no, LC.02.0076 | **901***, 902, **739**, **740**, 903, **923**, **924**, **741**, **925**, **721**, 742, **722** |
| | Avenacoside B (Aos_B) | yes, LC.03.0061 | **1247**, **1248**, **1249**, **1309**, 1278, 797, 1310, 1279, 650, 651, 1250, 1263 | no, LC.02.0441 | **1247**, **1248**, **1249**, **1309** |
| Avenacins | Avenacin A-1 (AEC_A1.1) | yes, LC.03.0066 | **1094**, **638**, **1095**, **152**, **639**, **1096***, 134, **1116**, 1117, **932**, **640**, **620** | no, LC.02.0099 | **638**, **1094**, 536, **1095**, **639**, **1096***, 537, **640**, **620**, **1116**, **932**, **152** |
| | Avenacin A-1 (AEC_A1.2) | yes, LC.03.0609 | **1094**, 1095, 638, 1096*, 639 | yes, LC.02.0551 | 98, **1094** |

**Table S2** ANOVA results for days to heading covariate significance in drBLUP calculation for metabolite and seed size phenotypes, where p-values meeting a bonferroni multiple test correction cutoff per panel and location are bolded. Seed length and width (and thus volume and surface area) are not available from the elite panel evaluated in South Dakota.

| | Diversity Panel | Elite panel, MN | Elite panel, SD | Elite panel, WI |
|---|---|---|---|---|
| AVN_A | F(1,362)=1.58, p=0.21 | F(1,208)=6.04, p=0.01 | F(1,215)=4.97, p=0.03 | F(1,209)=0, p=0.99 |
| AVN_B | F(1,364)=0.75, p=0.39 | F(1,211)=4.63, p=0.03 | F(1,216)=4.00, p=0.05 | F(1,211)=0.38, p=0.54 |
| AEC_A1.1 | F(1,364)=4.45, p=0.04 | F(1,223)=0.03, p=0.87 | F(1,207)=13.04, **p=3.5e-04** | F(1,210)=2.89, p=0.09 |
| AEC_A1.2 | F(1,364)=1.46, p=0.23 | F(1,215)=0.04, p=0.83 | F(1,210)=13.89, **p=2.49e-04** | F(1,207)=0.49, p=0.49 |
| AOS_A | F(1,362)=33.66, **p=1.44e-08** | F(1,214)=0.90, p=0.34 | F(1,218)=16.16, **p=8.00e-05** | F(1,226)=3.96, p=0.05 |
| AOS_dA | F(1,360)=5.36, p=0.02 | F(1,204)=0.90, p=0.34 | F(1,215)=6.79, p=0.01 | F(1,224)=26.3, **p=6.2e-07** |
| AOS_B | F(1,363)=11.46, **p=7.89e-04** | F(1,212)=3.45, p=0.06 | F(1,210)=0.41, p=0.52 | F(1,221)=3.64, p=0.06 |
| Seed volume | F(1,351)=0, p=0.98 | F(1,212)=12.87, **p=4.15e-04** | NA | F(1,198)=10.33, **p=1.53e-03** |
| Seed surface area | F(1,354)=3.22, p=0.07 | F(1,213)=16.97, **p=5.43e-05** | NA | F(1,200)=4.47, p=0.04 |
| Hundred kernel weight | F(1,366)=0.20, p=0.66 | F(1,214)=4.96, **p=0.03** | F(1,208)=29.77, **p=1.37e-07** | F(1,198)=22.58, **p=3.86e-06** |
| Hundred hull weight | F(1,362)=0.50, p=0.48 | F(1,210)=5.07, **p=0.03** | F(1,208)=0, p=1 | F(1,197)=0, p=0.95 |
| Percent groat | F(1,360)=0.38, p=0.54 | F(1,219)=0.82, p=0.82 | F(1,214)=28.65, **p=2.23e-07** | F(1,201)=21.36, **p=6.80e-06** |

**Table S3** Year of variety release from all available oat lines in diversity panel and source.

| T3 Name | Synonyms | Year | Source |
|---|---|---|---|
| TERRY | BGMN206; PI_25850 | 1909 | GRIN |
| WHITE_TARTARIAN | BGMN363; CI_800 | 1916 | GRIN |
| OAC_NO72 | CI_846 | 1919 | GRIN |
| GOLDEN_RUSTPROOF | BGMN346; CI_1751 | 1920 | GRIN |
| RED_TEXAS | BGMN156; CI_1914 | 1920 | GRIN |
| CORNELLIAN | SELECTION110_36 | 1920 | GRIN |
| KINVARRA_NO_A-8 | BGMN354; PI_306411 | 1928 | GRIN |
| BIHARIA | BGMN294; CI_5641 | 1933 | GRIN |
| TENNESSEE_SELECTION090 | BGMN116; CI_3175 | 1934 | GRIN |
| LENROC | | 1935 | https://doi.org/10.2134/agronj1935.00021962002700120007x |
| AMES_SELECTION4103 | BGMN102; CI_3328 | 1937 | GRIN |
| TIFT | | 1938 | GRIN |
| II-30-39 | BGMN479; CI_3655 | 1939 | GRIN |
| DUPPAWSKI | PI_131620 | 1939 | GRIN |
| LISCHOWER_FRUHHAFER | BGMN449; PI_178479; CIAV3799 | 1939 | GRIN |
| ABERDEEN_SELECTION1939-2005 | BGMN057; CI_3868 | 1940 | GRIN |
| ABERDEEN_SELECTION1939-3927 | BGMN051; PI_577978 | 1940 | GRIN |
| OSAGE | | 1941 | GRIN |
| CIAV4143 | BGMN170 | 1942 | GRIN |
| ANDREW | | 1942 | GRIN |
| MOHAWK | SELECTION1307_9 | 1943 | GRIN |
| FULGRAIN_SELECTION | BGMN258; CI_4565 | 1945 | GRIN |
| DELAIR | RESELECTION4076_16 | 1946 | GRIN |
| ABEGWEIT | BGMN492; CI_4970 | 1947 | GRIN |
| KHARKOVSKIJ596 | BGMN281 | 1947 | GRIN |
| PI159172 | BGMN109 | 1947 | GRIN |
| PI159180 | BGMN421 | 1947 | GRIN |
| FLORILAND | | 1947 | https://doi.org/10.2134/agronj1955.00021962004700110016x |
| MISSOURI04103 | BGMN555; CI_4987 | 1948 | GRIN |
| BRANCH | BGMN172; CI_5013 | 1948 | GRIN |

| | | | |
|---|---|---|---|
| CIAV5019 | BGMN193; CI_5019 | 1948 | GRIN |
| CD3708 | BGMN374; CI_5061 | 1948 | GRIN |
| CD3737 | BGMN371; CI_5068 | 1948 | GRIN |
| CD3774 | BGMN379; CI_5072 | 1948 | GRIN |
| PI168094 | BGMN569; | 1948 | GRIN |
| CIAV5033 | BGMN162; PI_290035 | 1948 | GRIN |
| CIAV5218 | BGMN590; PI_605532 | 1948 | GRIN |
| CIAV5220 | BGMN200; PI_605531 | 1948 | GRIN |
| CIAV5222 | BGMN272; CI_5220 | 1948 | GRIN |
| ENDRESS | BGMN103; PI_180920 | 1949 | GRIN |
| HOHENHEIMER_V | BGMN303; PI_180928 | 1949 | GRIN |
| LANG DOERFLERS WEIHENSTEPHANER WEISSHAFER | BGMN247; PI_180932 | 1949 | GRIN |
| CHLUMECKY | BGMN194; PI_182836 | 1949 | GRIN |
| CIAV5389 | CI_5218 | 1949 | GRIN |
| BLANCHE_DE_HONGRIE | BGMN001; CI_6246 | 1949 | GRIN |
| CIAV5666 | BGMN071 | 1950 | GRIN |
| CIAV5925 | BGMN130 | 1950 | GRIN |
| RANSOM | BGMN301; CI_5927 | 1950 | GRIN |
| CIAV5928 | BGMN364 | 1950 | GRIN |
| SELECTION3863-8 | BGMN573; CI_6034 | 1951 | GRIN |
| CIAV6130 | BGMN286 | 1951 | GRIN |
| CIAV6209 | | 1951 | GRIN |
| PI193957 | BGMN381 | 1951 | GRIN |
| PI194895 | | 1951 | GRIN |
| PI197400 | | 1951 | GRIN |
| SELECTION3841-1 | BGMN207; PI_605539 | 1951 | GRIN |
| CIAV6218 | BGMN383; CI_1642 | 1951 | GRIN |
| CIAV6227 | BGMN313; CI_3882 | 1951 | GRIN |
| CLINTLAND60 | INDIANA5413 | 1955 | https://doi.org/10.2134/agronj1958.00021962005000110022x |
| WIR4301 | PI_258641 | 1959 | GRIN |
| BENDERY878A\|PI258703 | BGMN150; PI_258703 | 1959 | GRIN |
| CLINTFORD | | 1959 | GRIN |
| TIOGA | 5217A1_2B_39 | 1959 | GRIN |
| NIAGARA | 5220A2_B_23 | 1959 | GRIN |
| WIR4672 | CI_6252 | 1959 | GRIN |
| TYLER | | 1961 | GRIN |
| TIPPECANOE | | 1961 | GRIN |

| | | | |
|---|---|---|---|
| ORBIT | 5279A1B_3B_70 | 1962 | GRIN |
| QUALITY\|PI289587 | PI_289587 | 1963 | GRIN |
| FLORIDA500 | PGR5886 | 1963 | GRIN |
| WIR4071 | BGMN093; PI_296150 | 1964 | GRIN |
| KRYMSKIJ90\|PI296174 | BGMN484; CI_6159 | 1964 | GRIN |
| ONOHOJSKIJ_A-547 | PI_326221 | 1968 | GRIN |
| SEVERNYJ209 | BGMN266; PI_326230 | 1968 | GRIN |
| SEVERIANIN\|PI326231 | BGMN087; PI_326231 | 1968 | GRIN |
| SKOROSPELKS | BGMN563; PI_326236 | 1968 | GRIN |
| Y-90 | BGMN410; PI_341021 | 1969 | GRIN |
| Y-498 | BGMN229; PI_341049 | 1969 | GRIN |
| PI344841 | PI_344841 | 1969 | GRIN |
| 26-35-B69 | BGMN003; PI_66489 | 1970 | GRIN |
| ARIANE\|PI361884 | BGMN262; PI_361884 | 1971 | GRIN |
| BANKUT_GALBEN | BGMN036; PI_361885 | 1971 | GRIN |
| BARAGAN114 | BGMN452; PI_361886 | 1971 | GRIN |
| CAUCAZ4275 | BGMN490; PI_361888 | 1971 | GRIN |
| CENAD88\|PI361889 | BGMN306; PI_361889 | 1971 | GRIN |
| DRUMMOND | | 1972 | https://triticeaetoolbox.org/POOL |
| HUSAR | BGMN359; PI_392043 | 1974 | GRIN |
| ALLEN | | 1974 | GRIN |
| VI2 | BGMN415; PI_504895 | 1974 | GRIN |
| AVOINE_NUE-NUE_NOISE | BGMN433; PI_401772 | 1975 | GRIN |
| STANTON | BGMN031; PI_412928 | 1976 | GRIN |
| LANG | | 1976 | GRIN |
| VI56 | BGMN341; PI_420442 | 1977 | GRIN |
| PI436081 | | 1979 | GRIN |
| PI436071 | BGMN366; PI_287294 | 1979 | GRIN |
| 77NZ_AA322 | BGMN117; PI_458807 | 1981 | GRIN |
| PENNLINE6571 | BGMN168; PI_469105 | 1982 | GRIN |
| SD751187 | BGMN328; PI_469265 | 1982 | GRIN |
| PA7733-1268 | BGMN285; PI_186618 | 1982 | GRIN |
| PERDEBERG | BGMN215; PI_479656 | 1983 | GRIN |
| FLAMINGSKOMET | BGMN468; PI_486323 | 1984 | GRIN |
| HAZEL | | 1985 | GRIN |
| PA7967-11759 | BGMN264; PI_504588 | 1986 | GRIN |
| MN861900 | BGMN067; PI_504954 | 1986 | GRIN |
| 2A-3 | BGMN352; PI_577858 | 1986 | GRIN |

| | | | |
|---|---|---|---|
| PI577862 | BGMN239 | 1986 | GRIN |
| 57A-3 | BGMN308; PI_577913 | 1986 | GRIN |
| 74C-1 | BGMN192; PI_577936 | 1986 | GRIN |
| 151C-2 | PI_577979 | 1986 | GRIN |
| PI577985 | BGMN163 | 1986 | GRIN |
| MN861218 | BGMN169; PI_290049 | 1986 | GRIN |
| KAPP | | 1986 | https://triticeaetoolbox.org/POOL |
| VALLEY | BGMN480; PI_525183 | 1988 | GRIN |
| MARION_QC | BGMN375; PI_536549 | 1989 | GRIN |
| PENNCOMP38 | BGMN041; PI_536614 | 1989 | GRIN |
| FLAEMINGSNOVA | | 1989 | https://triticeaetoolbox.org/POOL |
| NEWDAK | ND810104 | 1990 | GRIN |
| X397-1-B5-2 | BGMN394; PI_605547 | 1991 | GRIN |
| X345-1-B4-20-1 | BGMN076; PI_131621 | 1991 | GRIN |
| LENA | | 1994 | https://ec.europa.eu/food/plant/plant_propagation_material/plant_variety_catalogues_databases |
| JERRY | ND870952 | 1995 | GRIN |
| RODGERS | NC88_1818 | 1996 | GRIN |
| GRANE | | 1996 | https://triticeaetoolbox.org/POOL |
| BIRI | | 1997 | https://triticeaetoolbox.org/POOL |
| REEVES | SD97525 | 1997 | https://triticeaetoolbox.org/POOL |
| SECRETARIAT_LA495 | | 1999 | https://apps.ams.usda.gov/ |
| SYLVA | | 1999 | https://triticeaetoolbox.org/POOL |
| FREDDY | | 1999 | https://triticeaetoolbox.org/POOL |
| RANCH | | 1999 | https://triticeaetoolbox.org/POOL |
| LA604 | | 2000 | https://apps.ams.usda.gov/ |
| LUTZ | | 2000 | https://ec.europa.eu/food/plant/plant_propagation_material/plant_variety_catalogues_databases |
| RONALD | | 2000 | https://www.inspection.gc.ca/english/plaveg/pbrpov/cropreport/oat/ |
| BELINDA | | 2001 | https://ec.europa.eu/food/plant/plant_propagation_material/plant_variety_catalogues_databases |
| MAIDA | ND010264 | 2001 | https://triticeaetoolbox.org/POOL |
| CDC_ORRIN | | 2002 | https://www.inspection.gc.ca/english/plaveg/pbrpov/cropreport/oat/ |
| KOLBU | | 2003 | https://ec.europa.eu/food/plant/plant_propagation_material/plant_variety_catalogues_databases |
| SHERWOOD | OA1019_1, OA019_1 | 2004 | https://triticeaetoolbox.org/POOL |
| CDC_WEAVER | | 2005 | https://www.inspection.gc.ca/english/plaveg/pbrpov/cropreport/oat/ |
| CDC_SOL-FI | | 2005 | https://www.inspection.gc.ca/english/plaveg/pbrpov/cropreport/oat/ |
| IL2294-8 | | 2006 | GRIN |

| | | | |
|---|---|---|---|
| ROBUST | P973A38_9_3_27 | 2006 | https://triticeaetoolbox.org/POOL |
| HORIZON270 | | 2008 | https://apps.ams.usda.gov/ |
| NES | | 2008 | https://ec.europa.eu/food/plant/plant_propagation_material/plant_variety_catalogues_databases |
| CDC_MINSTREL | SO03191 | 2008 | https://www.inspection.gc.ca/english/plaveg/pbrpov/cropreport/oat/ |
| HORIZON201 | | 2009 | https://apps.ams.usda.gov/ |
| ODAL | | 2009 | https://ec.europa.eu/food/plant/plant_propagation_material/plant_variety_catalogues_databases |
| X8995-4 | | 2010 | GRIN |
| HY174-OA | OA1174_3 | 2010 | https://www.inspection.gc.ca/english/plaveg/pbrpov/cropreport/oat/ |
| CDC_SEABISCUIT | OT3036 | 2010 | https://www.inspection.gc.ca/english/plaveg/pbrpov/cropreport/oat/ |
| CORRAL | | 2011 | https://apps.ams.usda.gov/ |
| CDC_BIG_BROWN | OT3037 | 2011 | https://www.inspection.gc.ca/english/plaveg/pbrpov/cropreport/oat/ |
| BETAGENE | X8787-1 | 2012 | https://apps.ams.usda.gov/ |
| CILLA | | 2012 | https://ec.europa.eu/food/plant/plant_propagation_material/plant_variety_catalogues_databases |
| OPTIMUM | OA1228_1 | 2012 | https://www.inspection.gc.ca/english/plaveg/pbrpov/cropreport/oat/ |
| AAC_ROSKENS | OA1250_1 | 2012 | https://www.inspection.gc.ca/english/plaveg/pbrpov/cropreport/oat/ |

**Table S4** Crown rust QTL SNPs from recent publications mapped the most recent genome available (PepsiCO OT3098v1; https://wheat.pw.usda.gov/GG3/graingenes_downloads/oat-ot3098-pepsico).

| SNP | Chr | Pos | Source |
|---|---|---|---|
| GMI_DS_LB_10834 | 4A | 172124235 | Babiker *et al.*, 2015 |
| GMI_ES02_c27120_208 | 4A | 187629339 | Babiker *et al.*, 2015 |
| GMI_ES03_c6181_441 | 4A | 205590466 | Babiker *et al.*, 2015 |
| GMI_ES15_lrc9062_227 | 4A | 221497473 | Babiker *et al.*, 2015 |
| GMI_ES02_c6122_167 | 4A | 409045181 | Babiker *et al.*, 2015 |
| GMI_ES14_c1439_83 | 5D | 481192333 | Lin *et al.*, 2014 |
| GMI_GBS_90753 | 5D | 481977813 | Lin *et al.*, 2014 |
| GMI_ES03_c2277_336 | 1A | 388428416 | McNish *et al.*, 2020 |
| avgbs_cluster_33848.1.36 | 1A | 490650549 | McNish *et al.*, 2020 |
| avgbs_cluster_33848.1.37 | 1A | 490650550 | McNish *et al.*, 2020 |
| GMI_DS_LB_2908 | 1A | 511872969 | McNish *et al.*, 2020 |
| avgbs_200593.1.40 | 2A | 355735423 | McNish *et al.*, 2020 |
| GMI_ES03_c7453_413 | 2A | 443104084 | McNish *et al.*, 2020 |
| GMI_ES05_c11155_383 | 2D | 487988274 | McNish *et al.*, 2020 |
| avgbs_63606.1.44 | 3C | 595727680 | McNish *et al.*, 2020 |
| avgbs_cluster_10965.1.20 | 3C | 630246900 | McNish *et al.*, 2020 |
| avgbs2_56943.1.20 | 3C | 630246900 | McNish *et al.*, 2020 |
| GMI_ES13_c626_111 | 4A | 164054723 | McNish *et al.*, 2020 |
| avgbs_51923.1.27 | 4A | 268366966 | McNish *et al.*, 2020 |
| avgbs_87322.1.32 | 4A | 274622562 | McNish *et al.*, 2020 |
| GMI_ES02_c14986_166 | 4D | 240005580 | McNish *et al.*, 2020 |
| avgbs_120698.1.15 | 6A | 4281011 | McNish *et al.*, 2020 |
| GMI_DS_LB_270 | 6A | 428713105 | McNish *et al.*, 2020 |
| avgbs_cluster_15624.1.33 | 7A | 6403711 | McNish *et al.*, 2020 |
| avgbs_cluster_15624.1.63 | 7A | 6403741 | McNish *et al.*, 2020 |
| GMI_ES01_c27692_191 | 7A | 33516801 | McNish *et al.*, 2020 |
| GMI_ES22_c11318_631 | 7A | 43423184 | McNish *et al.*, 2020 |
| GMI_ES05_c14988_223 | 7A | 280368979 | McNish *et al.*, 2020 |
| GMI_ES18_c3370_505 | 7A | 347815005 | McNish *et al.*, 2020 |
| GMI_DS_A3_37_143 | 7A | 372943457 | McNish *et al.*, 2020 |
| GMI_ES17_c3808_324 | 7A | 376544507 | McNish *et al.*, 2020 |
| MI_ES17_c1629_493 | 7A | 394266638 | McNish *et al.*, 2020 |
| GMI_ES01_lrc22746_326 | 7A | 414338149 | McNish *et al.*, 2020 |

| | | | |
|---|---|---|---|
| avgbs2_85675.2.52 | 7D | 464421571 | McNish *et al.*, 2020 |
| avgbs_95303.1.18 | 7D | 507959223 | McNish *et al.*, 2020 |
| GMI_DS_LB_6017 | 4A | 450213722 | Zhao *et al.*, 2020 |
| GMI_ES14_lrc18344_662 | 4D | 376903720 | Zhao *et al.*, 2020 |

**Table S5** Transcripts associated with the avenanthramide biosynthetic and upstream shikimate pathway identified using Ensemble Enzyme Prediction Pipeline (E2P2) annotations.

| Transcript ID | Pathway Part | MetaCYC category |
|---|---|---|
| TRINITY_DN23077_c0_g2_i1 | phenylpropanoid | Phenylalanine-Ammonia-Lyase-RXN |
| TRINITY_DN26560_c0_g2_i1 | phenylpropanoid | Phenylalanine-Ammonia-Lyase-RXN |
| TRINITY_DN29757_c0_g1_i8 | phenylpropanoid | Phenylalanine-Ammonia-Lyase-RXN |
| TRINITY_DN32486_c0_g1_i5 | phenylpropanoid | Phenylalanine-Ammonia-Lyase-RXN |
| TRINITY_DN57700_c0_g1_i1 | phenylpropanoid | Phenylalanine-Ammonia-Lyase-RXN |
| TRINITY_DN12109_c0_g1_i1 | phenylpropanoid | Trans-Cinnamate-4-Monooxygenase-RXN |
| TRINITY_DN1934_c0_g1_i2 | phenylpropanoid | Trans-Cinnamate-4-Monooxygenase-RXN |
| TRINITY_DN1934_c1_g1_i1 | phenylpropanoid | Trans-Cinnamate-4-Monooxygenase-RXN |
| TRINITY_DN2359_c0_g1_i4 | phenylpropanoid | Trans-Cinnamate-4-Monooxygenase-RXN |
| TRINITY_DN10089_c0_g1_i4 | phenylpropanoid | 4-Coumarate--CoA-Ligase-RXN |
| TRINITY_DN1121_c1_g1_i2 | phenylpropanoid | 4-Coumarate--CoA-Ligase-RXN |
| TRINITY_DN1963_c0_g1_i1 | phenylpropanoid | 4-Coumarate--CoA-Ligase-RXN |
| TRINITY_DN22415_c0_g1_i4 | phenylpropanoid | 4-Coumarate--CoA-Ligase-RXN |
| TRINITY_DN26405_c0_g1_i5 | phenylpropanoid | 4-Coumarate--CoA-Ligase-RXN |
| TRINITY_DN29123_c0_g1_i1 | phenylpropanoid | 4-Coumarate--CoA-Ligase-RXN |
| TRINITY_DN37894_c0_g1_i1 | phenylpropanoid | RXN-2581 (CYP98A3) |
| TRINITY_DN6473_c0_g1_i1 | phenylpropanoid | RXN-2581 (CYP98A3) |
| TRINITY_DN23025_c0_g1_i1 | phenylpropanoid | 2.3.1.133-RXN (HHCT) |
| TRINITY_DN26554_c0_g1_i3 | phenylpropanoid | 2.3.1.133-RXN (HHCT) |
| TRINITY_DN5172_c0_g1_i4 | phenylpropanoid | 2.3.1.133-RXN (HHCT) |
| TRINITY_DN13049_c0_g2_i4 | phenylpropanoid | RXN-2621 (HHCT2) |
| TRINITY_DN14241_c0_g1_i1 | phenylpropanoid | RXN-2621 (HHCT2) |
| TRINITY_DN31721_c0_g1_i6 | phenylpropanoid | RXN-2621 (HHCT2) |
| TRINITY_DN46541_c0_g1_i2 | phenylpropanoid | RXN-2621 (HHCT2) |
| TRINITY_DN56875_c0_g1_i1 | phenylpropanoid | RXN-2621 (HHCT2) |
| TRINITY_DN3508_c0_g1_i7 | phenylpropanoid | NA, identified as CCoAOMT in Hu (2020) |
| TRINITY_DN1249_c1_g1_i2 | shikimate | DAHPSYN-RXN |
| TRINITY_DN3009_c0_g1_i1 | shikimate | DAHPSYN-RXN |
| TRINITY_DN5456_c0_g1_i1 | shikimate | DAHPSYN-RXN |
| TRINITY_DN8688_c0_g1_i2 | shikimate | 3-Dehydroquinate-Synthase-RXN |
| TRINITY_DN1649_c0_g1_i2 | shikimate | 3-Dehydroquinate-Dehydratase-RXN |
| TRINITY_DN2726_c0_g1_i2 | shikimate | 3-Dehydroquinate-Dehydratase-RXN |
| TRINITY_DN1649_c0_g1_i2 | shikimate | Shikimate-5-Dehydrogenase -RXN |
| TRINITY_DN2726_c0_g1_i2 | shikimate | Shikimate-5-Dehydrogenase-RXN |
| TRINITY_DN10783_c0_g2_i1 | shikimate | Shikimate-Kinase-RXN |
| TRINITY_DN10783_c0_g3_i1 | shikimate | Shikimate-Kinase-RXN |
| TRINITY_DN3895_c0_g1_i7 | shikimate | Shikimate-Kinase-RXN |

| | | |
|---|---|---|
| TRINITY_DN1033_c0_g1_i2 | shikimate | Chorismate-Synthase-RXN |
| TRINITY_DN1661_c0_g1_i1 | anthranilate synthase | ANTHRANSYN-RXN |

**Table S6** Heritability of measured seed size parameters (length, width, height, hundred kernel weight, hundred hull weight and groat percent) and calculated values (volume, surface area, and surface area to volume ratio). Seed length and width (and thus volume and surface area) are not available from the elite panel evaluated in South Dakota.

| $h^2$ | Diversity panel | Elite panel, MN | Elite panel, SD | Elite panel, WI |
|---|---|---|---|---|
| Seed length | 0.53 | 0.66 | NA | 0.44 |
| Seed width | 0.67 | 0.62 | NA | 0.45 |
| Seed height | 0.68 | 0.29 | 0.44 | 0.44 |
| Seed volume | 0.72 | 0.50 | NA | 0.33 |
| Seed surface area | 0.65 | 0.57 | NA | 0.32 |
| Seed SA:Volume ratio | 0.69 | 0.25 | NA | 0.38 |
| Hundred kernel weight | 0.79 | 0.45 | 0.49 | 0.26 |
| Hundred hull weight | 0.57 | 0.26 | 0.19 | 0.00 |
| Groat percent | 0.62 | 0.79 | 0.27 | 0.58 |

**Table S7** Relationship between seed size and metabolite abundance ANOVA results, where significant results are bolded.

| Panel | Seed volume | Seed surface area | Surface area to volume ratio |
|---|---|---|---|
| *Diverse* | | | |
| AVN_A | F(1,347)=50.84, **p=5.85e-12** | F(1,345)=43.84, **p=1.37e-10** | F(1,348)=41.49, **p=3.95e-10** |
| AVN_B | F(1,348)=54.28, **p=1.27e-12** | F(1,346)=48.57, **p=1.62e-11** | F(1,349)=41.15, **p=4.59e-10** |
| AEC_A1.1 | F(1,345)=0.29, p=0.59 | F(1,343)=0.02, p=0.89 | F(1,346)=1.04, p=0.31 |
| AEC_A1.2 | F(1,346)=0.05, p=0.82 | F(1,344)=0.03, p=0.87 | F(1,347)=0.74, p=0.39 |
| AOS_A | F(1,346)=62.62, **p=3.43e-14** | F(1,344)=46.49, **p=4.31e-11** | F(1,347)=74.16, **p=2.56e-16** |
| AOS_dA | F(1,345)=36.90, **p=3.29e-09** | F(1,343)=28.13, **p=2.04e-07** | F(1,346)=39.15, **p=1.16e-09** |
| AOS_B | F(1,347)=34.00, **p=1.26e-08** | F(1,345)=25.12, **p=8.61e-07** | F(1,348)=30.62, **p=6.19e-08** |
| *Elite, MN* | | | |
| AVN_A | F(1,205)=15.64, **p=1.04e-04** | F(1,205)=14.46, **p=1.89e-04** | F(1,205)=11.89, **p=6.85e-04** |
| AVN_B | F(1,208)=13.69, **p=2.75e-04** | F(1,208)=10.57, **p=1.34e-03** | F(1,207)=10.35, **p=1.51e-03** |
| AEC_A1.1 | F(1,209)=1.11, p=0.29 | F(1,209)=1.71, p=0.19 | F(1,208)=0.59, p=0.44 |
| AEC_A1.2 | F(1,208)=0.33, p=0.56 | F(1,208)=0.66, p=0.42 | F(1,207)=0.06, p=0.80 |
| AOS_A | F(1,209)=2.04, p=0.15 | F(1,209)=1.34, p=0.25 | F(1,208)=3.22, p=0.07 |
| AOS_dA | F(1,209)=3.49, p=0.06 | F(1,209)=2.13, p=0.15 | F(1,208)=4.27, **p=0.04** |
| AOS_B | F(1,208)=1.41, p=0.24 | F(1,208)=0.72, p=0.40 | F(1,207)=1.91, p=0.17 |
| *Elite, SD* | | | |
| | NA | NA | NA |
| *Elite, WI* | | | |
| AVN_A | F(1,194)=24.33, **p=1.74e-06** | F(1,197)=29.12, **p=1.94e-07** | F(1,194)=13.53, **p=3.03e-04** |
| AVN_B | F(1,193)=20.37, **p=1.11e-05** | F(1,196)=24.86, **p=1.34e-06** | F(1,193)=12.21, **p=5.90e-04** |
| AEC_A1.1 | F(1,195)=17.25, **p=4.90e-05** | F(1,198)=16.36, **p=7.49e-05** | F(1,195)=15.26, **p=1.29e-04** |
| AEC_A1.2 | F(1,195)=12.60, **p=4.83e-04** | F(1,198)=13.09, **p=3.76e-04** | F(1,195)=7.15, **p=8.13e-03** |

**Table S8** Relationship between seed weight and metabolite abundance ANOVA results, where significant results are bolded.

| Panel | Hundred Kernel Weight | Hundred Hull Weight | Percent groat |
|---|---|---|---|
| *Diverse* | | | |
| AVN_A | F(1,359)=42.77, **p=2.12e-10** | F(1,354)=13.38, **p=2.93e-04** | F(1,356)=23.86, **p=1.57e-06** |
| AVN_B | F(1,360)=52.58, **p=2.55e-12** | F(1,355)=14.02, **p=2.11e-04** | F(1,357)=26.55, **p=4.26e-07** |
| AEC_A1.1 | F(1,357)=0.24, p=0.62 | F(1,352)=0.07, p=0.79 | F(1,354)=0.73, p=0.39 |
| AEC_A1.2 | F(1,358)=0.00, p=0.97 | F(1,353)=0.41, p=0.52 | F(1,355)=0.01, p=0.91 |
| AOS_A | F(1,358)=55.35, **p=7.54e-13** | F(1,353)=26.16, **p=5.17e-07** | F(1,355)=4.28, **p=0.04** |
| AOS_dA | F(1,357)=38.96, **p=1.23e-09** | F(1,352)=18.13, **p=2.66e-05** | F(1,354)=5.64, **p=0.02** |
| AOS_B | F(1,359)=37.46, **p=2.44e-09** | F(1,354)=11.79, **p=6.66e-04** | F(1,356)=6.31, **p=0.01** |
| *Elite, MN* | | | |
| AVN_A | F(1,206)=10.71, **p=1.25e-03** | F(1,204)=1.31, p=0.25 | F(1,204)=1.83, p=0.09 |
| AVN_B | F(1,209)=12.59, **p=4.79e-04** | F(1,207)=2.38, p=0.12 | F(1,207)=1.61, p=0.21 |
| AEC_A1.1 | F(1,210)=0.26, p=0.61 | F(1,208)=1.31, p=0.25 | F(1,208)=0.90, p=0.34 |
| AEC_A1.2 | F(1,209)=0.25, p=0.62 | F(1,207)=0.68, p=0.41 | F(1,207)=0.40, p=0.53 |
| AOS_A | F(1,210)=0.98, p=0.32 | F(1,208)=1.22, p=0.27 | F(1,208)=0.04, p=0.84 |
| AOS_dA | F(1,210)=1.80, p=0.18 | F(1,208)=0.73, p=0.39 | F(1,208)=0.16, p=0.69 |
| AOS_B | F(1,209)=1.84, p=0.18 | F(1,207)=0.47, p=0.50 | F(1,207)=0.14, p=0.71 |
| *Elite, SD* | | | |
| AVN_A | F(1,200)=29.51, **p=1.61e-07** | F(1,203)=16.90, **p=5.73e-05** | F(1,200)=0.27, p=0.60 |
| AVN_B | F(1,200)=30.06, **p=1.26e-07** | F(1,203)=13.26, **p=3.44e-04** | F(1,200)=1.17, p=0.28 |
| AEC_A1.1 | F(1,199)=1.12, p=0.29 | F(1,202)=3.54, p=0.06 | F(1,199)=1.11, p=0.29 |
| AEC_A1.2 | F(1,199)=0.10, p=0.76 | F(1,202)=1.45, p=0.23 | F(1,199)=1.07, p=0.30 |
| AOS_A | F(1,197)=6.53, **p=0.01** | F(1,200)=0.25, p=0.62 | F(1,197)=4.11, **p=0.04** |
| AOS_dA | F(1,199)=2.21, p=0.14 | F(1,202)=0.64, p=0.43 | F(1,199)=5.75, **p=0.02** |
| AOS_B | F(1,200)=0.92, p=0.34 | F(1,203)=0.01, p=0.92 | F(1,200)=1.02, p=0.31 |
| *Elite, WI* | | | |
| AVN_A | F(1,197)=22.56, **p=3.92e-06** | F(1,198)=0.15, p=0.70 | F(1,195)=20.91, **p=8.54e-06** |

| | | | |
|---|---|---|---|
| AVN_B | F(1,196)=19.78, **p=1.45e-05** | F(1,197)=0.75, p=0.39 | F(1,194)=14.76, **p=1.65e-04** |
| AEC_A1.1 | F(1,198)=9.95, **p=1.86e-03** | F(1,199)=0.82, p=0.37 | F(1,196)=5.67, **p=0.02** |
| AEC_A1.2 | F(1,197)=7.29, **p=7.53e-03** | F(1,198)=0.12, p=0.73 | F(1,195)=6.46, **p=0.01** |

**Table S9** Genetic correlation between seed size phenotypes (seed volume, “seedVol”; seed surface area, “seedSA”; hundred kernel weight, “HKW”; and groat percent, “groatPct”) and specialized metabolites by panel. The number of samples in each panel are indicated. Seed surface area and volume phenotypes were not available from South Dakota.

| | AVN_A | AVN_B | AEC_A1.1 | AEC_A1.2 | AOS_A | AOS_dA | AOS_B |
|---|---|---|---|---|---|---|---|
| *Diversity* | | | | | | | |
| seedVol | 0.70 | 0.71 | -0.04 | -0.04 | -0.30 | -0.28 | -0.31 |
| seedSA | 0.71 | 0.74 | -0.01 | 0.00 | -0.23 | -0.24 | -0.25 |
| HKW | 0.74 | 0.88 | -0.13 | -0.12 | -0.46 | -0.39 | -0.45 |
| groatPct | 0.25 | 0.43 | -0.26 | -0.16 | -0.27 | -0.02 | -0.40 |
| *Elite, MN* | | | | | | | |
| seedVol | 0.03 | -0.29 | -0.12 | -0.08 | -0.51 | -0.74 | NA |
| seedSA | 0.07 | -0.31 | -0.09 | -0.07 | -0.43 | -0.66 | NA |
| HKW | 0.57 | 0.23 | 0.02 | 0.09 | -0.19 | -0.51 | NA |
| groatPct | -0.17 | -0.17 | -0.44 | -0.46 | 0.02 | -0.15 | NA |
| *Elite, SD* | | | | | | | |
| seedVol | NA | NA | NA | NA | NA | NA | NA |
| seedSA | NA | NA | NA | NA | NA | NA | NA |
| HKW | 0.54 | 0.70 | 0.24 | -0.26 | -0.45 | NA | NA |
| groatPct | 0.34 | 0.37 | -0.08 | 0.08 | -0.20 | NA | NA |
| *Elite, WI* | | | | | | | |

| | | | | | | | |
|---|---|---|---|---|---|---|---|
| seedVol | 0.58 | 0.15 | 0.15 | 0.08 | NA | NA | NA |
| seedSA | 0.70 | 0.22 | 0.14 | 0.10 | NA | NA | NA |
| HKW | 0.79 | 0.57 | -0.03 | 0.04 | NA | NA | NA |
| groatPct | 0.77 | 0.51 | 0.10 | 0.00 | NA | NA | NA |

**Table S10** Relationships between variety release year and seed size and metabolite abundance

ANOVA results, where significant results are bolded.

| Phenotype | Variety release year |
|---|---|
| Seed volume | F(1,138)=26.32, **p=9.63e-07** |
| Seed surface area | F(1,137)=18.00, **p=4.13e-05** |
| Seed surface area to volume ratio | F(1,139)=25.00, **p=1.70e-06** |
| Hundred kernel weight | F(1,145)=19.21, **p=2.23e-05** |
| Hundred hull weight | F(1,143)=17.25, **p=5.62e-05** |
| Percent groat | F(1,144)= 0.00, p=0.95 |
| AVN_A | F(1,143)= 1.53, p=0.22 |
| AVN_B | F(1,144)=0.19, p=0.67 |
| AEC_A1.1 | F(1,143)=3.11, p=0.08 |
| AEC_A1.2 | F(1,143)=5.75, **p=0.02** |
| AOS_A | F(1,144)=8.55, **p=4.02e-03** |
| AOS_dA | F(1,142)=7.58, **p=6.67e-03** |
| AOS_B | F(1,144)=3.40, p=0.07 |

**Table S11** Results from multiple regression analysis using variety release year and seed size. The model parameters presented are degrees of freedom and adjusted $R^2$ value. For both predictors, the coefficient with standard error and the *p*-value that the coefficient is significantly different from zero are presented. Bolded values indicate *p*<0.006 to account for bonferroni adjusted multiple test corrections per trait.

| Phenotype | Model parameters | Variety release year | Seed volume |
|---|---|---|---|
| Percent groat | df = 133; $R^2_{adj}$= 0.09 | ß= -0.017 +/- 0.010; P(>\|t\|) = 0.09 | **ß= 0.225 +/- 0.058; P(>\|t\|) < 0.001** |
| AVN_A | df = 133; $R^2_{adj}$= 0.05 | ß= -0.001 +/- 0.003; P(>\|t\|) = 0.78 | **ß= 0.055 +/- 0.019; P(>\|t\|) = 0.004** |
| AVN_B | df = 134; $R^2_{adj}$= 0.04 | ß= -0.005 +/- 0.005; P(>\|t\|) = 0.31 | ß= 0.072 +/- 0.026; P(>\|t\|) = 0.006 |
| AEC_A1.1 | df = 133; $R^2_{adj}$= 0.01 | ß= -0.005 +/- 0.003; P(>\|t\|) = 0.16 | ß= 0.027 +/- 0.019; P(>\|t\|) = 0.16 |
| AEC_A1.2 | df = 133; $R^2_{adj}$= 0.02 | ß= -0.007 +/- 0.003; P(>\|t\|) = 0.04 | ß= 0.026 +/- 0.019; P(>\|t\|) = 0.18 |
| AOS_A | df = 134; $R^2_{adj}$= 0.14 | ß= -0.001 +/- 0.001; P(>\|t\|) = 0.33 | **ß= -0.016 +/- 0.004; P(>\|t\|) < 0.001** |
| AOS_dA | df = 132; $R^2_{adj}$= 0.05 | ß= -0.002 +/- 0.002; P(>\|t\|) = 0.23 | **ß= -0.024 +/- 0.008; P(>\|t\|) = 0.005** |
| AOS_B | df = 134; $R^2_{adj}$= 0.02 | ß= -0.001 +/- 0.008; P(>\|t\|) = 0.25 | ß= -0.005 +/- 0.005; P(>\|t\|) = 0.23 |

**Table S12** All transcripts within 100kb of significant GWAS results. The distance from the start and end of the transcript are given in kb where negative corresponds to upstream and positive is downstream.

| Transcript ID | Chr | start | end | Distance from GWAS SNP (kB) |
|---|---|---|---|---|
| TRINITY_DN8881_c0_g1_i7[1] | 2D | 518526065 | 518531763 | (-38.3, -44) |
| TRINITY_DN82490_c0_g1_i1 | 2D | 518401587 | 518401913 | (86.2, 85.9) |
| TRINITY_DN27158_c0_g1_i1[2] | 2D | 518393691 | 518398259 | (94.1, 89.5) |
| TRINITY_DN10810_c1_g1_i1 | 3A | 406969353 | 406972256 | (59.8, 62.7) |
| TRINITY_DN107261_c0_g1_i1 | 3C | 3658370 | 3658621 | (-3.7, -4.0) |
| TRINITY_DN108057_c0_g1_i1 | 3C | 3656805 | 3657438 | (-2.2, -2.8) |
| TRINITY_DN113235_c0_g1_i1 | 3C | 3561553 | 3561772 | (93.1, 92.9) |
| TRINITY_DN19987_c0_g2_i1 | 3C | 3659197 | 3660196 | (-4.6, -5.6) |
| TRINITY_DN30319_c0_g2_i2 | 3C | 3656414 | 3656662 | (-1.8, -2.0) |
| TRINITY_DN38516_c0_g1_i1 | 3C | 3657443 | 3658185 | (-2.8, -3.5) |
| TRINITY_DN43715_c0_g1_i1 | 3C | 3652092 | 3652407 | (2.6, 2.2) |
| TRINITY_DN4392_c2_g1_i1 | 3C | 3661439 | 3661798 | (-6.8, -7.2) |
| TRINITY_DN70089_c0_g1_i1 | 3C | 3658599 | 3659055 | (-4.0, -4.4) |
| TRINITY_DN89608_c0_g1_i1 | 3C | 3653387 | 3653657 | (1.3, 1.0) |
| TRINITY_DN95686_c0_g1_i1 | 3C | 3561111 | 3561314 | (93.5, 93.3) |
| TRINITY_DN41508_c0_g2_i1 | 3C | 6151753 | 6153225 | (49.7, 48.2) |
| TRINITY_DN41508_c0_g4_i1 | 3C | 6153202 | 6153430 | (48.3, 48.0) |
| TRINITY_DN4832_c0_g1_i3[3] | 3C | 6184681 | 6188160 | (16.8, 13.3) |
| TRINITY_DN68354_c0_g1_i1 | 3C | 6153439 | 6153742 | (48.0, 47.7) |
| TRINITY_DN96860_c0_g1_i1 | 3C | 6187161 | 6187421 | (14.3, 14.0) |
| TRINITY_DN102553_c0_g1_i1 | 3C | 7251691 | 7251936 | (41.5, 41.3) |
| TRINITY_DN109320_c0_g1_i1 | 3C | 7249030 | 7249275 | (44.2, 43.9) |
| TRINITY_DN110193_c0_g1_i1 | 3C | 7314970 | 7315545 | (-21.8, -22.3) |
| TRINITY_DN20068_c0_g1_i4[4] | 3C | 7242798 | 7254266 | (50.4, 38.9) |
| TRINITY_DN40977_c0_g1_i1 | 3C | 7243511 | 7244055 | (49.7, 49.2) |
| TRINITY_DN51708_c0_g1_i1 | 3C | 7247612 | 7247825 | (45.6, 45.4) |
| TRINITY_DN75514_c0_g1_i1 | 3C | 7247108 | 7247452 | (46.1, 45.8) |
| TRINITY_DN76174_c0_g1_i1 | 3C | 7220732 | 7221148 | (72.5, 72.1) |
| TRINITY_DN76297_c0_g1_i1 | 3C | 7254689 | 7255029 | (38.5, 38.2) |
| TRINITY_DN76434_c0_g1_i1 | 3C | 7223129 | 7223352 | (70.1, 69.9) |
| TRINITY_DN83830_c0_g1_i1 | 3C | 7222305 | 7222544 | (70.9, 70.7) |
| TRINITY_DN85096_c0_g1_i1 | 3C | 7222911 | 7223114 | (70.3, 70.1) |
| TRINITY_DN9136_c1_g2_i1 | 4D | 266011904 | 266012419 | (-83.3, -82.8) |
| TRINITY_DN43775_c0_g1_i1[5] | 4D | 266037947 | 266039657 | (-57.2, -55.5) |
| TRINITY_DN110558_c0_g1_i1 | 4D | 266061567 | 266061889 | (-33.6, -33.3) |
| TRINITY_DN115400_c0_g1_i1 | 4D | 266062972 | 266063198 | (-32.2, -32.0) |

| | | | | |
|---|---|---|---|---|
| TRINITY_DN258_c0_g1_i4 | 4D | 266067188 | 266068310 | (-28.0, -26.9) |
| TRINITY_DN12531_c0_g1_i1 | 4D | 266093259 | 266150492 | (-1.9, 55.3) |
| TRINITY_DN84630_c0_g1_i1 | 4D | 266094550 | 266094835 | (-0.6, -0.4) |
| TRINITY_DN9136_c1_g3_i1 | 4D | 266143371 | 266144045 | (48.2, 48.9) |
| TRINITY_DN51300_c0_g1_i1 | 4D | 266150569 | 266151627 | (55.4, 56.4) |
| TRINITY_DN4758_c0_g1_i2 | 4D | 266151576 | 266152330 | (56.4, 57.1) |
| TRINITY_DN38906_c0_g1_i1 | 4D | 266152095 | 266152768 | (56.9, 57.6) |
| TRINITY_DN23092_c2_g1_i1 | 5A | 456590215 | 456591682 | (89.2, 90.7) |
| TRINITY_DN111982_c0_g1_i1 | 5D | 387417550 | 387417810 | (40.6, 40.9) |
| TRINITY_DN102666_c0_g1_i1 | 5D | 387418886 | 387419292 | (42.0, 42.4) |
| TRINITY_DN30583_c0_g1_i1 | 5D | 387420951 | 387421994 | (44.0, 45.1) |
| TRINITY_DN57375_c0_g1_i2 | 5D | 387422796 | 387423134 | (45.9, 46.2) |

[1]Annotated asO-fucosyltransferase 15
[2]Annotated asZinc finger BED domain-containing protein RICESLEEPER 2
[3]Annotated as L-Ala-D/L-amino acid epimerase
[4]Annotated as LEAF RUST 10 DISEASE-RESISTANCE LOCUS RECEPTOR-LIKE PROTEIN KINASE-like 1.2
[5]Annotated as UDP-glycosyltransferase 91D1

**Table S13** Full avenanthramide transcriptome-wide association study (TWAS) results of all transcripts with $p_{FDR}$<0.05. Trait indicates the avenanthramides, and the original and FDR corrected p-value, and effect and standard error of effect are provided.

| trait | transcript | Rank_by_trait | pValue | effect | effectSe | pfdr |
|---|---|---|---|---|---|---|
| LC.03.0125 | TRINITY_DN8445_c0_g1_i4 | 1 | 9.10E-10 | 0.575 | 0.097 | 1.03E-05 |
| LC.03.0125 | TRINITY_DN21184_c0_g2_i1 | 2 | 8.00E-10 | 0.444 | 0.074 | 1.03E-05 |
| LC.03.0125 | TRINITY_DN8412_c0_g1_i1 | 3 | 4.80E-08 | 0.423 | 0.079 | 0.0002 |
| LC.03.0125 | TRINITY_DN826_c0_g1_i8 | 4 | 4.77E-08 | 0.677 | 0.127 | 0.0002 |
| LC.03.0125 | TRINITY_DN21184_c0_g4_i1 | 5 | 2.93E-08 | 0.399 | 0.074 | 0.0002 |
| LC.03.0125 | TRINITY_DN13908_c0_g4_i1 | 6 | 7.72E-08 | 0.455 | 0.087 | 0.0002 |
| LC.03.0125 | TRINITY_DN3664_c1_g1_i1 | 7 | 7.52E-08 | 0.455 | 0.086 | 0.0002 |
| LC.03.0125 | TRINITY_DN3664_c0_g3_i1 | 8 | 1.90E-07 | 0.497 | 0.097 | 0.0005 |
| LC.03.0125 | TRINITY_DN22256_c0_g1_i3 | 9 | 7.79E-07 | 0.408 | 0.084 | 0.0015 |
| LC.03.0125 | TRINITY_DN60211_c0_g1_i1 | 10 | 6.68E-07 | 0.700 | 0.143 | 0.0015 |
| LC.03.0125 | TRINITY_DN11606_c0_g1_i4 | 11 | 6.77E-07 | 0.619 | 0.127 | 0.0015 |
| LC.03.0125 | TRINITY_DN68113_c0_g1_i1 | 12 | 7.47E-07 | 0.530 | 0.109 | 0.0015 |
| LC.03.0125 | TRINITY_DN784_c0_g1_i3 | 13 | 1.21E-06 | 0.817 | 0.171 | 0.0021 |
| LC.03.0125 | TRINITY_DN8304_c0_g1_i4 | 14 | 2.48E-06 | 0.696 | 0.150 | 0.0037 |
| LC.03.0125 | TRINITY_DN2924_c0_g1_i2 | 15 | 2.56E-06 | 0.987 | 0.213 | 0.0037 |
| LC.03.0125 | TRINITY_DN10815_c0_g1_i1 | 16 | 2.60E-06 | 0.623 | 0.135 | 0.0037 |
| LC.03.0125 | TRINITY_DN3664_c0_g1_i5 | 17 | 3.05E-06 | 0.699 | 0.152 | 0.0038 |
| LC.03.0125 | TRINITY_DN4109_c0_g1_i3 | 18 | 3.06E-06 | 0.953 | 0.207 | 0.0038 |
| LC.03.0125 | TRINITY_DN1008_c0_g2_i2 | 19 | 3.59E-06 | 0.880 | 0.193 | 0.0043 |
| LC.03.0125 | TRINITY_DN39942_c0_g1_i1 | 20 | 4.04E-06 | 1.066 | 0.235 | 0.0046 |
| LC.03.0125 | TRINITY_DN26560_c0_g2_i1 | 21 | 4.35E-06 | 0.274 | 0.061 | 0.0047 |
| LC.03.0125 | TRINITY_DN1581_c0_g1_i3 | 22 | 4.82E-06 | 1.004 | 0.223 | 0.0050 |
| LC.03.0125 | TRINITY_DN43_c0_g1_i3 | 23 | 5.86E-06 | 0.278 | 0.062 | 0.0058 |
| LC.03.0125 | TRINITY_DN2877_c0_g1_i6 | 24 | 7.75E-06 | 0.417 | 0.095 | 0.0062 |
| LC.03.0125 | TRINITY_DN21916_c0_g1_i1 | 25 | 7.94E-06 | 0.510 | 0.116 | 0.0062 |
| LC.03.0125 | TRINITY_DN29096_c0_g1_i9 | 26 | 7.99E-06 | 0.879 | 0.200 | 0.0062 |
| LC.03.0125 | TRINITY_DN26866_c0_g2_i3 | 27 | 6.87E-06 | 0.271 | 0.061 | 0.0062 |
| LC.03.0125 | TRINITY_DN13998_c0_g2_i1 | 28 | 7.71E-06 | 0.490 | 0.111 | 0.0062 |
| LC.03.0125 | TRINITY_DN15065_c0_g2_i1 | 29 | 7.06E-06 | 0.487 | 0.110 | 0.0062 |
| LC.03.0125 | TRINITY_DN16295_c0_g1_i1 | 30 | 9.84E-06 | 0.520 | 0.119 | 0.0074 |
| LC.03.0125 | TRINITY_DN14541_c0_g1_i1 | 31 | 1.03E-05 | 0.396 | 0.091 | 0.0075 |
| LC.03.0125 | TRINITY_DN1103_c0_g1_i1 | 32 | 1.07E-05 | 0.976 | 0.225 | 0.0075 |
| LC.03.0125 | TRINITY_DN2577_c0_g1_i1 | 33 | 1.10E-05 | 0.781 | 0.180 | 0.0076 |
| LC.03.0125 | TRINITY_DN15878_c0_g1_i6 | 34 | 1.19E-05 | 0.462 | 0.107 | 0.0077 |
| LC.03.0125 | TRINITY_DN28530_c0_g1_i4 | 35 | 1.18E-05 | 0.824 | 0.191 | 0.0077 |
| LC.03.0125 | TRINITY_DN21184_c0_g3_i1 | 36 | 1.60E-05 | 0.469 | 0.110 | 0.0095 |
| LC.03.0125 | TRINITY_DN16348_c0_g1_i1 | 37 | 1.64E-05 | -0.691 | 0.163 | 0.0095 |
| LC.03.0125 | TRINITY_DN19061_c0_g1_i1 | 38 | 1.62E-05 | 0.741 | 0.174 | 0.0095 |
| LC.03.0125 | TRINITY_DN3916_c0_g1_i1 | 39 | 1.52E-05 | 0.718 | 0.168 | 0.0095 |
| LC.03.0125 | TRINITY_DN43068_c0_g1_i1 | 40 | 2.07E-05 | 0.382 | 0.091 | 0.0117 |
| LC.03.0125 | TRINITY_DN11938_c0_g1_i1 | 41 | 2.35E-05 | 0.548 | 0.131 | 0.0130 |

| | | | | | | |
|---|---|---|---|---|---|---|
| LC.03.0125 | TRINITY_DN5351_c0_g1_i2 | 42 | 2.45E-05 | 0.590 | 0.141 | 0.0132 |
| LC.03.0125 | TRINITY_DN4266_c0_g1_i6 | 43 | 2.55E-05 | 0.736 | 0.177 | 0.0133 |
| LC.03.0125 | TRINITY_DN91049_c0_g1_i1 | 44 | 2.64E-05 | 0.517 | 0.124 | 0.0133 |
| LC.03.0125 | TRINITY_DN337_c0_g1_i1 | 45 | 2.64E-05 | 1.025 | 0.247 | 0.0133 |
| LC.03.0125 | TRINITY_DN49524_c0_g1_i1 | 46 | 2.95E-05 | 0.484 | 0.117 | 0.0141 |
| LC.03.0125 | TRINITY_DN42793_c0_g2_i1 | 47 | 2.98E-05 | 0.348 | 0.084 | 0.0141 |
| LC.03.0125 | TRINITY_DN8115_c0_g1_i2 | 48 | 2.96E-05 | 0.790 | 0.191 | 0.0141 |
| LC.03.0125 | TRINITY_DN1291_c0_g1_i1 | 49 | 3.16E-05 | 0.886 | 0.215 | 0.0146 |
| LC.03.0125 | TRINITY_DN78280_c0_g1_i1 | 50 | 3.49E-05 | 0.349 | 0.085 | 0.0158 |
| LC.03.0125 | TRINITY_DN13413_c0_g1_i2 | 51 | 3.95E-05 | 0.733 | 0.180 | 0.0175 |
| LC.03.0125 | TRINITY_DN276_c1_g1_i1 | 52 | 4.09E-05 | 0.403 | 0.099 | 0.0178 |
| LC.03.0125 | TRINITY_DN2744_c0_g1_i4 | 53 | 4.58E-05 | 0.762 | 0.189 | 0.0196 |
| LC.03.0125 | TRINITY_DN23931_c0_g1_i6 | 54 | 5.13E-05 | 0.587 | 0.147 | 0.0215 |
| LC.03.0125 | TRINITY_DN18727_c0_g2_i1 | 55 | 5.34E-05 | 0.557 | 0.139 | 0.0220 |
| LC.03.0125 | TRINITY_DN13684_c0_g1_i1 | 56 | 5.71E-05 | 0.806 | 0.203 | 0.0231 |
| LC.03.0125 | TRINITY_DN3411_c0_g1_i4 | 57 | 5.96E-05 | 0.621 | 0.156 | 0.0236 |
| LC.03.0125 | TRINITY_DN2351_c0_g1_i1 | 58 | 6.05E-05 | -1.312 | 0.331 | 0.0236 |
| LC.03.0125 | TRINITY_DN8268_c1_g1_i3 | 59 | 6.96E-05 | -0.581 | 0.147 | 0.0267 |
| LC.03.0125 | TRINITY_DN13177_c0_g1_i1 | 60 | 7.19E-05 | 0.635 | 0.162 | 0.0271 |
| LC.03.0125 | TRINITY_DN2385_c0_g1_i1 | 61 | 8.48E-05 | 0.729 | 0.187 | 0.0310 |
| LC.03.0125 | TRINITY_DN2768_c0_g1_i2 | 62 | 8.38E-05 | 0.818 | 0.210 | 0.0310 |
| LC.03.0125 | TRINITY_DN5540_c0_g1_i2 | 63 | 8.93E-05 | -1.126 | 0.290 | 0.0321 |
| LC.03.0125 | TRINITY_DN6507_c0_g1_i4 | 64 | 0.000100267 | 0.340 | 0.088 | 0.0355 |
| LC.03.0125 | TRINITY_DN8686_c0_g3_i1 | 65 | 0.000102555 | 0.849 | 0.221 | 0.0357 |
| LC.03.0125 | TRINITY_DN2667_c0_g1_i1 | 66 | 0.000120934 | 0.741 | 0.195 | 0.0415 |
| LC.03.0125 | TRINITY_DN1363_c0_g1_i2 | 67 | 0.00012934 | -1.005 | 0.265 | 0.0437 |
| LC.03.0125 | TRINITY_DN2212_c0_g1_i2 | 68 | 0.000131947 | -1.228 | 0.324 | 0.0439 |
| LC.03.0125 | TRINITY_DN112536_c0_g1_i1 | 69 | 0.000135805 | -0.427 | 0.113 | 0.0446 |
| LC.03.0125 | TRINITY_DN19306_c0_g1_i4 | 70 | 0.000141883 | 0.410 | 0.109 | 0.0459 |
| LC.03.0125 | TRINITY_DN1191_c0_g1_i15 | 71 | 0.000148804 | -0.927 | 0.247 | 0.0474 |
| LC.03.0125 | TRINITY_DN42577_c0_g1_i1 | 72 | 0.000154948 | 0.371 | 0.099 | 0.0487 |
| LC.03.1204 | TRINITY_DN8445_c0_g1_i4 | 1 | 7.29E-11 | 0.476 | 0.075 | 9.21E-07 |
| LC.03.1204 | TRINITY_DN21184_c0_g2_i1 | 2 | 8.14E-11 | 0.366 | 0.058 | 9.21E-07 |
| LC.03.1204 | TRINITY_DN21184_c0_g4_i1 | 3 | 4.74E-09 | 0.328 | 0.057 | 3.57E-05 |
| LC.03.1204 | TRINITY_DN8412_c0_g1_i1 | 4 | 8.38E-09 | 0.348 | 0.062 | 4.74E-05 |
| LC.03.1204 | TRINITY_DN3664_c0_g3_i1 | 5 | 5.11E-08 | 0.405 | 0.076 | 0.0002 |
| LC.03.1204 | TRINITY_DN826_c0_g1_i8 | 6 | 8.07E-08 | 0.520 | 0.099 | 0.0003 |
| LC.03.1204 | TRINITY_DN4109_c0_g1_i3 | 7 | 1.29E-07 | 0.837 | 0.162 | 0.0004 |
| LC.03.1204 | TRINITY_DN68113_c0_g1_i1 | 8 | 1.66E-07 | 0.437 | 0.085 | 0.0005 |
| LC.03.1204 | TRINITY_DN13998_c0_g2_i1 | 9 | 2.17E-07 | 0.441 | 0.087 | 0.0005 |
| LC.03.1204 | TRINITY_DN11606_c0_g1_i4 | 10 | 1.96E-07 | 0.506 | 0.099 | 0.0005 |
| LC.03.1204 | TRINITY_DN13908_c0_g4_i1 | 11 | 2.79E-07 | 0.341 | 0.068 | 0.0006 |
| LC.03.1204 | TRINITY_DN26866_c0_g2_i3 | 12 | 3.69E-07 | 0.238 | 0.048 | 0.0007 |
| LC.03.1204 | TRINITY_DN3664_c1_g1_i1 | 13 | 4.15E-07 | 0.335 | 0.067 | 0.0007 |
| LC.03.1204 | TRINITY_DN22256_c0_g1_i3 | 14 | 8.31E-07 | 0.318 | 0.066 | 0.0013 |
| LC.03.1204 | TRINITY_DN1008_c0_g2_i2 | 15 | 1.01E-06 | 0.724 | 0.151 | 0.0015 |
| LC.03.1204 | TRINITY_DN15065_c0_g2_i1 | 16 | 1.20E-06 | 0.410 | 0.086 | 0.0017 |
| LC.03.1204 | TRINITY_DN15878_c0_g1_i6 | 17 | 1.35E-06 | 0.397 | 0.084 | 0.0018 |
| LC.03.1204 | TRINITY_DN14541_c0_g1_i1 | 18 | 1.47E-06 | 0.336 | 0.071 | 0.0018 |

| | | | | | | |
|---|---|---|---|---|---|---|
| LC.03.1204 | TRINITY_DN60211_c0_g1_i1 | 19 | 1.81E-06 | 0.526 | 0.112 | 0.0022 |
| LC.03.1204 | TRINITY_DN3664_c0_g1_i5 | 20 | 3.16E-06 | 0.545 | 0.119 | 0.0034 |
| LC.03.1204 | TRINITY_DN26560_c0_g2_i1 | 21 | 3.25E-06 | 0.217 | 0.047 | 0.0034 |
| LC.03.1204 | TRINITY_DN1103_c0_g1_i1 | 22 | 3.26E-06 | 0.804 | 0.175 | 0.0034 |
| LC.03.1204 | TRINITY_DN2744_c0_g1_i4 | 23 | 3.99E-06 | 0.671 | 0.148 | 0.0038 |
| LC.03.1204 | TRINITY_DN49524_c0_g1_i1 | 24 | 3.98E-06 | 0.416 | 0.092 | 0.0038 |
| LC.03.1204 | TRINITY_DN10815_c0_g1_i1 | 25 | 4.62E-06 | 0.475 | 0.105 | 0.0042 |
| LC.03.1204 | TRINITY_DN29096_c0_g1_i9 | 26 | 7.12E-06 | 0.690 | 0.156 | 0.0062 |
| LC.03.1204 | TRINITY_DN6507_c0_g1_i4 | 27 | 7.86E-06 | 0.304 | 0.069 | 0.0066 |
| LC.03.1204 | TRINITY_DN11938_c0_g1_i1 | 28 | 8.25E-06 | 0.451 | 0.103 | 0.0067 |
| LC.03.1204 | TRINITY_DN12758_c2_g2_i1 | 29 | 1.00E-05 | 0.389 | 0.089 | 0.0078 |
| LC.03.1204 | TRINITY_DN512_c0_g2_i1 | 30 | 1.11E-05 | 0.542 | 0.125 | 0.0079 |
| LC.03.1204 | TRINITY_DN2577_c0_g1_i1 | 31 | 1.10E-05 | 0.610 | 0.141 | 0.0079 |
| LC.03.1204 | TRINITY_DN276_c1_g1_i1 | 32 | 1.12E-05 | 0.337 | 0.078 | 0.0079 |
| LC.03.1204 | TRINITY_DN8304_c0_g1_i4 | 33 | 1.27E-05 | 0.505 | 0.117 | 0.0087 |
| LC.03.1204 | TRINITY_DN4595_c0_g1_i8 | 34 | 1.31E-05 | 0.555 | 0.129 | 0.0087 |
| LC.03.1204 | TRINITY_DN23931_c0_g1_i6 | 35 | 1.50E-05 | 0.489 | 0.114 | 0.0094 |
| LC.03.1204 | TRINITY_DN43068_c0_g1_i1 | 36 | 1.49E-05 | 0.303 | 0.071 | 0.0094 |
| LC.03.1204 | TRINITY_DN91049_c0_g1_i1 | 37 | 1.88E-05 | 0.411 | 0.097 | 0.0115 |
| LC.03.1204 | TRINITY_DN3411_c0_g1_i4 | 38 | 2.49E-05 | 0.508 | 0.122 | 0.0146 |
| LC.03.1204 | TRINITY_DN13998_c0_g1_i1 | 39 | 2.58E-05 | 0.291 | 0.070 | 0.0146 |
| LC.03.1204 | TRINITY_DN13413_c0_g1_i2 | 40 | 2.52E-05 | 0.586 | 0.141 | 0.0146 |
| LC.03.1204 | TRINITY_DN16295_c0_g1_i1 | 41 | 2.68E-05 | 0.386 | 0.093 | 0.0148 |
| LC.03.1204 | TRINITY_DN2877_c0_g1_i6 | 42 | 2.97E-05 | 0.305 | 0.074 | 0.0160 |
| LC.03.1204 | TRINITY_DN21916_c0_g1_i1 | 43 | 3.29E-05 | 0.371 | 0.090 | 0.0173 |
| LC.03.1204 | TRINITY_DN21184_c0_g3_i1 | 44 | 4.06E-05 | 0.349 | 0.086 | 0.0209 |
| LC.03.1204 | TRINITY_DN1272_c0_g1_i3 | 45 | 4.17E-05 | 0.609 | 0.150 | 0.0210 |
| LC.03.1204 | TRINITY_DN43_c0_g1_i3 | 46 | 4.33E-05 | 0.196 | 0.049 | 0.0213 |
| LC.03.1204 | TRINITY_DN3916_c0_g1_i1 | 47 | 4.97E-05 | 0.527 | 0.131 | 0.0232 |
| LC.03.1204 | TRINITY_DN1581_c0_g1_i3 | 48 | 5.01E-05 | 0.698 | 0.174 | 0.0232 |
| LC.03.1204 | TRINITY_DN42577_c0_g1_i1 | 49 | 4.98E-05 | 0.310 | 0.077 | 0.0232 |
| LC.03.1204 | TRINITY_DN14735_c0_g2_i1 | 50 | 5.41E-05 | -0.327 | 0.082 | 0.0245 |
| LC.03.1204 | TRINITY_DN5351_c0_g1_i2 | 51 | 5.65E-05 | 0.440 | 0.110 | 0.0251 |
| LC.03.1204 | TRINITY_DN1405_c0_g1_i1 | 52 | 5.97E-05 | 0.396 | 0.100 | 0.0260 |
| LC.03.1204 | TRINITY_DN3267_c0_g1_i1 | 53 | 6.36E-05 | -0.642 | 0.162 | 0.0269 |
| LC.03.1204 | TRINITY_DN14356_c1_g1_i10 | 54 | 6.53E-05 | 0.528 | 0.134 | 0.0269 |
| LC.03.1204 | TRINITY_DN784_c0_g1_i3 | 55 | 6.49E-05 | 0.528 | 0.134 | 0.0269 |
| LC.03.1204 | TRINITY_DN27969_c0_g2_i1 | 56 | 7.03E-05 | 0.257 | 0.065 | 0.0284 |
| LC.03.1204 | TRINITY_DN39942_c0_g1_i1 | 57 | 7.33E-05 | 0.720 | 0.183 | 0.0291 |
| LC.03.1204 | TRINITY_DN11233_c0_g1_i7 | 58 | 7.69E-05 | 0.628 | 0.161 | 0.0295 |
| LC.03.1204 | TRINITY_DN1403_c1_g1_i1 | 59 | 7.62E-05 | 0.370 | 0.095 | 0.0295 |
| LC.03.1204 | TRINITY_DN16348_c0_g1_i1 | 60 | 8.75E-05 | -0.493 | 0.127 | 0.0326 |
| LC.03.1204 | TRINITY_DN16663_c0_g1_i1 | 61 | 8.78E-05 | 0.354 | 0.091 | 0.0326 |
| LC.03.1204 | TRINITY_DN20857_c0_g1_i4 | 62 | 9.69E-05 | 0.346 | 0.090 | 0.0352 |
| LC.03.1204 | TRINITY_DN27969_c0_g1_i1 | 63 | 9.80E-05 | 0.297 | 0.077 | 0.0352 |
| LC.03.1204 | TRINITY_DN12960_c1_g1_i1 | 64 | 0.000101623 | 0.274 | 0.071 | 0.0359 |
| LC.03.1204 | TRINITY_DN4096_c0_g1_i1 | 65 | 0.000103708 | -1.062 | 0.276 | 0.0361 |
| LC.03.1204 | TRINITY_DN2924_c0_g1_i2 | 66 | 0.000114934 | 0.636 | 0.166 | 0.0394 |
| LC.03.1204 | TRINITY_DN13889_c1_g1_i1 | 67 | 0.000130681 | -0.444 | 0.117 | 0.0442 |

| | | | | | | |
|---|---|---|---|---|---|---|
| LC.03.1204 | TRINITY_DN626_c0_g2_i3 | 68 | 0.000146766 | 0.493 | 0.131 | 0.0472 |
| LC.03.1204 | TRINITY_DN112536_c0_g1_i1 | 69 | 0.000146501 | -0.332 | 0.088 | 0.0472 |
| LC.03.1204 | TRINITY_DN13684_c0_g1_i1 | 70 | 0.000142335 | 0.596 | 0.158 | 0.0472 |
| LC.03.1204 | TRINITY_DN9961_c0_g1_i7 | 71 | 0.000148 | 0.375 | 0.100 | 0.0472 |
| LC.03.1204 | TRINITY_DN7337_c0_g3_i1 | 72 | 0.000158526 | 0.323 | 0.086 | 0.0498 |

**Table S14** The top three GO enrichment of biological process terms from avenacin (AEC) TWAS results. The *p*-values are not adjusted, and no terms are significant after false discovery rate adjustment.

| GO.ID | Term | AEC_A1.1 | | AEC_A1.2 | |
|---|---|---|---|---|---|
| | | rank | *p* | rank | *p* |
| GO:0009742 | brassinosteroid mediated signaling pathway | 1 | 0.006 | | |
| GO:0034727 | piecemeal microautophagy of the nucleus | 2 | 0.007 | 1 | 0.002 |
| GO:0015977 | carbon fixation | 3 | 0.010 | | |
| GO:0051028 | mRNA transport | 7 | 0.022 | 2 | 0.005 |
| GO:0045899 | positive regulation of RNA polymerase II | 9 | 0.023 | 3 | 0.010 |

**Table S15** The top three GO enrichment of biological process terms from avenacoside (AOS) TWAS results. The *p*-values are not adjusted, and no terms are significant after false discovery rate adjustment.

| GO.ID | Term | AOS_A | | AOS_dA | | AOS_B | |
|---|---|---|---|---|---|---|---|
| | | rank | *p* | rank | *p* | rank | *p* |
| GO:0009688 | abscisic acid biosynthetic process | 1 | 0.002 | - | - | - | - |
| GO:0010027 | thylakoid membrane organization | 2 | 0.006 | 2 | 0.004 | - | - |
| GO:0006289 | nucleotide-excision repair | 3 | 0.006 | - | - | - | - |
| GO:0010206 | photosystem II repair | 16 | 0.017 | 1 | 0.001 | - | - |
| GO:0006097 | glyoxylate cycle | - | - | 3 | 0.009 | - | - |
| GO:0016560 | protein import into peroxisome matrix | - | - | - | - | 1 | 0.001 |
| GO:0006281 | DNA repair | - | - | - | - | 2 | 0.008 |
| GO:0010228 | vegetative to reproductive phase | - | - | - | - | 3 | 0.009 |

**Table S16** The coefficient of determination between expression of all TWAS results shared by all avenanthramides ($p_{FDR}$ <0.05) and avenanthramide B ("$R^2_{adj}$_AVN_B") and between seed volume and TWAS gene expression ("$R^2_{adj}$_SeedVol"). The difference is presented where positive values indicate that the coefficient of determination for AVN_B was greater than that of seed volume.

| Transcript | $R^2_{adj}$_AVN_B | $R^2_{adj}$_SeedVol | Difference |
|---|---|---|---|
| TRINITY_DN8445_c0_g1_i4 | 0.129 | 0.012 | 0.117 |
| TRINITY_DN21184_c0_g2_i1 | 0.118 | 0.004 | 0.114 |
| TRINITY_DN826_c0_g1_i8 | 0.124 | 0.016 | 0.108 |
| TRINITY_DN784_c0_g1_i3 | 0.104 | 0.001 | 0.103 |
| TRINITY_DN3664_c1_g1_i1 | 0.097 | -0.002 | 0.099 |
| TRINITY_DN60211_c0_g1_i1 | 0.101 | 0.002 | 0.099 |
| TRINITY_DN11606_c0_g1_i4 | 0.102 | 0.006 | 0.097 |
| TRINITY_DN8412_c0_g1_i1 | 0.119 | 0.025 | 0.095 |
| TRINITY_DN3664_c0_g3_i1 | 0.116 | 0.022 | 0.094 |
| TRINITY_DN13908_c0_g4_i1 | 0.108 | 0.015 | 0.094 |
| TRINITY_DN21184_c0_g4_i1 | 0.094 | 0.013 | 0.081 |
| TRINITY_DN8304_c0_g1_i4 | 0.080 | 0.003 | 0.078 |
| TRINITY_DN26560_c0_g2_i1 | 0.087 | 0.009 | 0.078 |
| TRINITY_DN22256_c0_g1_i3 | 0.091 | 0.014 | 0.077 |
| TRINITY_DN1008_c0_g2_i2 | 0.081 | 0.004 | 0.076 |
| TRINITY_DN15878_c0_g1_i6 | 0.074 | -0.002 | 0.076 |
| TRINITY_DN16295_c0_g1_i1 | 0.079 | 0.004 | 0.075 |
| TRINITY_DN10815_c0_g1_i1 | 0.090 | 0.015 | 0.075 |
| TRINITY_DN3411_c0_g1_i4 | 0.081 | 0.007 | 0.074 |
| TRINITY_DN68113_c0_g1_i1 | 0.075 | 0.001 | 0.074 |
| TRINITY_DN15065_c0_g2_i1 | 0.087 | 0.013 | 0.074 |
| TRINITY_DN5351_c0_g1_i2 | 0.071 | -0.003 | 0.073 |
| TRINITY_DN39942_c0_g1_i1 | 0.091 | 0.019 | 0.073 |
| TRINITY_DN16348_c0_g1_i1 | 0.070 | -0.002 | 0.073 |
| TRINITY_DN2744_c0_g1_i4 | 0.078 | 0.007 | 0.071 |
| TRINITY_DN21916_c0_g1_i1 | 0.073 | 0.003 | 0.070 |
| TRINITY_DN91049_c0_g1_i1 | 0.070 | 0.002 | 0.068 |
| TRINITY_DN4109_c0_g1_i3 | 0.081 | 0.014 | 0.068 |
| TRINITY_DN2577_c0_g1_i1 | 0.069 | 0.002 | 0.067 |

| | | | |
|---|---|---|---|
| TRINITY_DN29096_c0_g1_i9 | 0.071 | 0.004 | 0.067 |
| TRINITY_DN43068_c0_g1_i1 | 0.074 | 0.011 | 0.064 |
| TRINITY_DN43_c0_g1_i3 | 0.074 | 0.010 | 0.064 |
| TRINITY_DN1581_c0_g1_i3 | 0.063 | 0.000 | 0.063 |
| TRINITY_DN2924_c0_g1_i2 | 0.074 | 0.011 | 0.063 |
| TRINITY_DN6507_c0_g1_i4 | 0.062 | 0.000 | 0.061 |
| TRINITY_DN11938_c0_g1_i1 | 0.067 | 0.007 | 0.060 |
| TRINITY_DN1103_c0_g1_i1 | 0.057 | 0.000 | 0.057 |
| TRINITY_DN14541_c0_g1_i1 | 0.068 | 0.011 | 0.057 |
| TRINITY_DN3916_c0_g1_i1 | 0.066 | 0.010 | 0.056 |
| TRINITY_DN21184_c0_g3_i1 | 0.058 | 0.003 | 0.055 |
| TRINITY_DN26866_c0_g2_i3 | 0.059 | 0.005 | 0.054 |
| TRINITY_DN3664_c0_g1_i5 | 0.057 | 0.003 | 0.054 |
| TRINITY_DN42577_c0_g1_i1 | 0.051 | -0.002 | 0.052 |
| TRINITY_DN276_c1_g1_i1 | 0.059 | 0.008 | 0.050 |
| TRINITY_DN49524_c0_g1_i1 | 0.068 | 0.018 | 0.050 |
| TRINITY_DN13413_c0_g1_i2 | 0.063 | 0.014 | 0.048 |
| TRINITY_DN13998_c0_g2_i1 | 0.073 | 0.028 | 0.045 |
| TRINITY_DN23931_c0_g1_i6 | 0.042 | -0.003 | 0.045 |
| TRINITY_DN2877_c0_g1_i6 | 0.077 | 0.033 | 0.044 |
| TRINITY_DN13684_c0_g1_i1 | 0.063 | 0.026 | 0.037 |
| TRINITY_DN112536_c0_g1_i1 | 0.024 | 0.004 | 0.020 |

**Methods S1** Metabolite extraction, measurement and annotation

Fifty mature seeds were dehulled, and the 100 mg samples were homogenized and extracted using a biphasic extraction method to separate polar and non-polar compounds using successive extractions beginning with with 33% methanol 67% methyl-tert-butyl-ether (MTBE) (V/V). The organic phase was removed, and then 50% methanol 50% acetonitrile (v/v) was added to precipitate proteins and excess glycans. The aqueous layer was then dried under nitrogen gas overnight and then re-suspended in 50% methanol 50% water (v/v) for further analysis. Chromatography analysis was done using a Waters Acquity UPLC system and compounds were separated using a Waters Acquity UPLC CSH Phenyl Hexyl column (1.7 μM, 1.0 x 100 mm), using a gradient from solvent A (2mM ammonium hydroxide, 0.1% formic acid) to solvent B (Acetonitrile, 0.1% formic acid). The column eluent was infused into a Waters Xevo G2 TOF-MS with an electrospray source in positive mode, scanning 50-2000 m/z at 0.2 seconds per scan, alternating between MS (6 V collision energy) and MSE mode (15-30 V ramp).

Raw data files were converted to .cdf format, and a matrix of molecular features as defined by retention time and mass (m/z) was generated using XCMS software in R (Smith *et al.*, 2006) for feature detection and alignment using the 'centWave' algorithm. Features were grouped using RAMClustR (Broeckling *et al.*, 2014), with normalization set to 'none'. LC-MS data were first annotated by searching against an in-house spectra and retention time database using RAMSearch (Broeckling *et al.*, 2016). RAMClustR was used to call the 'findMain' function from the interpretMSSpectrum (Jaeger *et al.*, 2017) package to infer the molecular weight of each LC-MS compound and annotate the mass signals. The complete MS spectrum and MSE spectrum were import to MSFinder (Tsugawa *et al.*, 2016) to determine the most probable molecular formula and structure and to perform a spectral search against the MassBank database. All results were imported into R and a collective annotation is derived with prioritization of RAMSearch > MSFinder mssearch > MSFinder structure > MSFinder formula > findMain M. Annotation confidence is reported as described (Sumner *et al.*, 2007).

Names and spectra of the specialized metabolites are given in **Table S1.** The mass spectra of the specialized metabolites were positively annotated by these methods in the diversity panel, which was analyzed in 2018. Many of the specialized metabolites were also annotated in the elite panel (measured in 2017), and missing annotations were completed by comparing spectra to the diversity panel and published mass spectra for avenanthramides (de Bruijn *et al.*, 2019), avenacins (Leveau *et al.*, 2019) and avenacosides (Bahraminejad *et al.*, 2008).
